# Pleiotropic and Epistatic Network-Based Discovery: Integrated Networks for Target Gene Discovery

**DOI:** 10.1101/267997

**Authors:** Deborah Weighill, Piet Jones, Manesh Shah, Priya Ranjan, Wellington Muchero, Jeremy Schmutz, Avinash Sreedasyam, David Macaya-Sanz, Robert Sykes, Nan Zhao, Madhavi Z. Martin, Stephen DiFazio, Timothy J. Tschaplinski, Gerald Tuskan, Daniel Jacobson

**Affiliations:** The Bredesen Center for Interdisciplinary Research and Graduate Education, University of Tennessee, Knoxville, Knoxville, TN, USA; Biosciences Division, Oak Ridge National Laboratory, Oak Ridge, TN, USA; National Renewable Energy Laboratory, Golden, CO, USA; Department of Energy Joint Genome Institute, Walnut Creek, CA, USA; HudsonAlpha Institute for Biotechnology, Huntsville, AL, USA; Department of Biology, West Virginia University, Morgantown, WV, USA; The University of Tennessee Institute of Agriculture, University of Tennessee, Knoxville, Knoxville, TN, USA

**Keywords:** *Multi-omic data layering*, *LOE Scores*, *Lines of Evidence Scores*, *GWAS*, *SNP correlation*, *networks*, *lignin*, *recalcitrance*, *bioenergy*, *co-expression*, *co-methylation*, *metabolomics*, *pyMBMS*

## Abstract

Biological organisms are complex systems that are composed of functional networks of interacting molecules and macromolecules. Complex phenotypes are the result of orchestrated, hierarchical, heterogeneous collections of expressed genomic variants. However, the effects of these variants are the result of historic selective pressure and current environmental and epigenetic signals, and, as such, their co-occurrence can be seen as genome-wide correlations in a number of different manners. Biomass recalcitrance (i.e., the resistance of plants to degradation or deconstruction, which ultimately enables access to a plant’s sugars) is a complex polygenic phenotype of high importance to biofuels initiatives. This study makes use of data derived from the re-sequenced genomes from over 800 different Populus trichocarpa genotypes in combination with metabolomic and pyMBMS data across this population, as well as co-expression and co-methylation networks in order to better understand the molecular interactions involved in recalcitrance, and identify target genes involved in lignin biosynthesis/degradation. A Lines Of Evidence (LOE) scoring system is developed to integrate the information in the different layers and quantify the number of lines of evidence linking genes to lignin-related lignin-phenotypes across the network layers. The resulting Genome Wide Association Study networks, integrated with Single Nucleotide Polymorphism (SNP) correlation, co-methylation and co-expression networks through the LOE scores are proving to be a powerful approach to determine the pleiotropic and epistatic relationships underlying cellular functions and, as such, the molecular basis for complex phenotypes, such as recalcitrance.

## 2. INTRODUCTION

*Populus* species are promising sources of cellulosic biomass for biofuels because of their fast growth rate, high cellulose content and moderate lignin content (Sannigrahi et al., 2010). Ragauskas et al. (2006) outline areas of research needed “to increase the impact, efficiency, and sustainability of bio-refinery facilities” (Ragauskas et al., 2006), such as research into modifying plants to enhance favorable traits, including altered cell wall structure leading to increased sugar release, as well as resilience to biotic and abiotic stress. One particular research target in *Populus* species is the decrease/alteration of the lignin content of cell walls.

A large collection of different data types has been generated for *Populus trichocarpa*. The genome has been sequenced and annotated (Tuskan et al., 2006), and the assembly is currently in its third version of revision. A collection of 1,100 accessions of *P. trichocarpa* that have been clonally propagated in four different common gardens (Tuskan et al., 2011; Slavov et al., 2012; Evans et al., 2014) have been resequenced, which has provided a large set of ~ 28,000,000 Single Nucleotide Polymorphisms (SNPs) that has recently been publicly released (http://bioenergycenter.org/besc/gwas/). Many molecular phenotypes, such as untargetted metabolomics and pyMBMS phenotypes, that have been measured in this population provide an unparalleled resource for Genome Wide Association Studies (for example, see McKown et al. (2014)). DNA methylation data in the form of MeDIP (Methyl-DNA immunoprecipitation)-seq has been performed on 10 different *P. trichocarpa* tissues (Vining et al., 2012), and gene expression has been measured across various tissues and conditions.

This study involves integrating these various data types in order to identify new possible candidate genes involved in lignin biosynthesis/degradation/regulation. Integrating Genome Wide Association Study (GWAS) data with other data types has previously been done to help provide context and identify relevant subnetworks/modules (Calabrese et al., 2017; Bunyavanich et al., 2014). Ritchie et al. (2015) reviewed techniques for integrating various data types for the aim of investigating gene-phenotype associations. Integrating multiple lines of evidence is a useful strategy as the more lines of evidence that connect a gene to a phenotype lowers the chance of false positives. Ritchie et al. (2015) categorized data integration approaches into two main classes, namely multi-staged analysis and meta-dimensional analysis. Multi-staged analysis analyses aims to enrich a biological signal through various steps of analysis. Meta-dimensional analysis involves the concurrent analysis of various data types, and is divided into three subcategories (Ritchie et al., 2015): Concatenation-based integration concatenates the data matrices of different data types into a single matrix on which a model is constructed (for example, see Fridley et al. (2012)). Model-based integration involves constructing a separate model for each dataset and then constructing a final model from the results of the separate models (for example, see Kim et al. (2013)). Transformation-based integration involves transforming transforming each data type into a common form (e.g. a network) before combining them (see for example, Kim et al. (2012)).

This study involves the development of an approach which can be seen as a type of transformation-based integration. Association networks for various different data types were constructed, including a pyMBMS GWAS network, a metabolomics GWAS network, as well as co-expression, co-methylation and SNP correlation networks, and subsequently the information in the different networks was integrated through the calculation of the newly developed Lines Of Evidence (LOE) scores defined in this study. These scores quantify the number of lines of evidence connecting each gene to lignin-related genes and phenotypes. This multi-omic data integration approach allowed for the identification of new possible candidate genes involved in lignin biosynthesis/regulation through multiple lines of evidence.

## 3. METHODS

### 3.1. Overview

This approach involved combining various data types in order to identify new possible target genes involved in lignin biosynthesis/degradation/regulation. Figure 1 summarizes the overall approach. First, association networks were constructed including metabolomics and pyMBMS GWAS networks, co-expression, co-methylation and SNP correlation networks. Known lignin-related genes and phenotypes were then identified, and used as seeds to select lignin-related subnetworks from these various networks. The Lines Of Evidence (LOE) scoring technique was developed, and each gene was then scored based on its Lines Of Evidence linking it to lignin-related genes and phenotypes.

**Figure 1:**
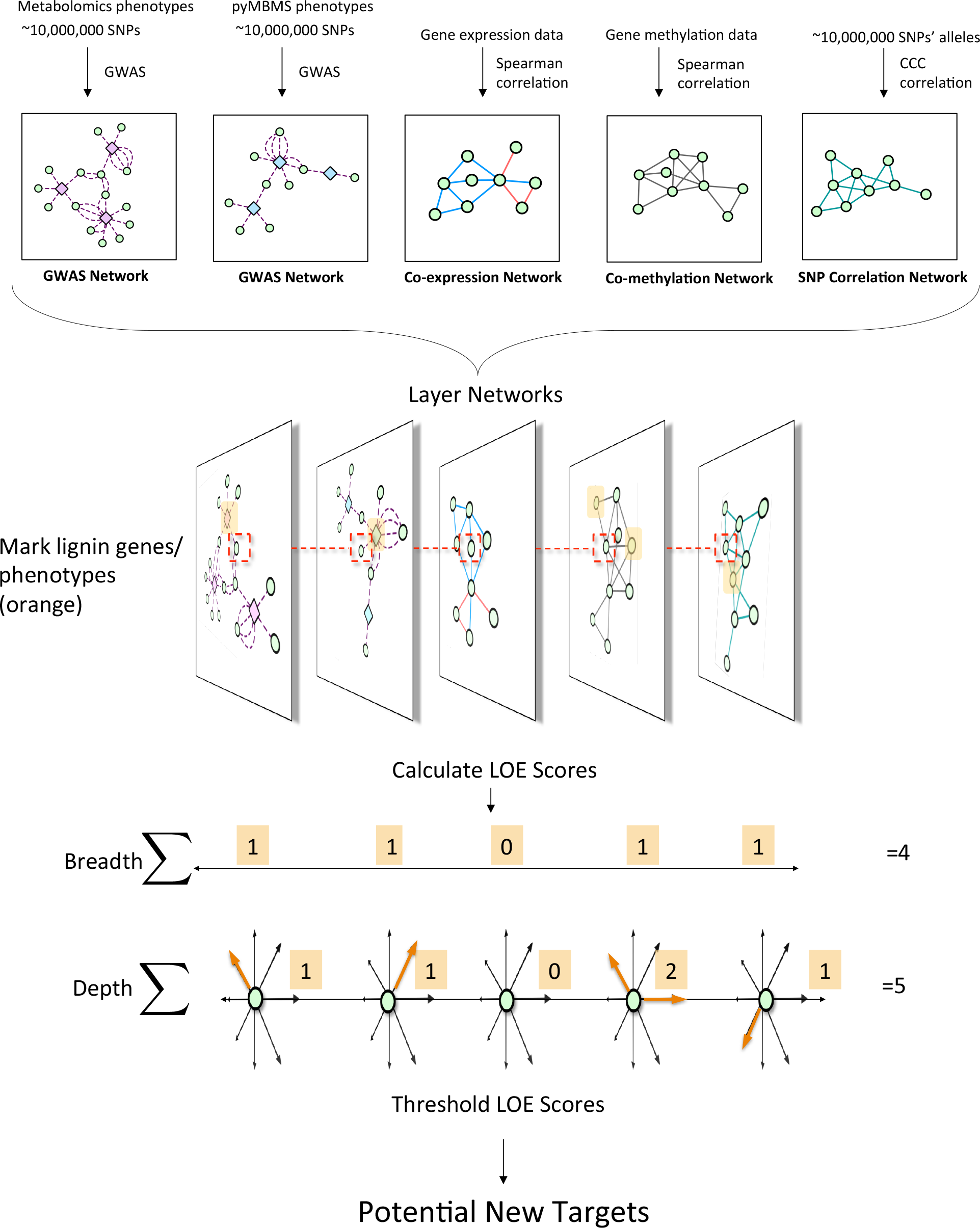
Overview of pipeline for data layering and score calcualtion. First, the different network layers are constructed. Networks are layered, and lignin-related genes and phenotypes (orange) are identified. LOE scores are calculated for each gene. An example of the LOE score calculation for the red-boxed gene is shown. Thresholding the LOE scores results in a set of new potential target genes involved in lignin biosynthesis/degradation/regulation.

### 3.2. GWAS Network Construction

#### 3.2.1 Metabolomics Data

The *P. trichocarpa* leaf samples for 851 unique clones were collected over three consecutive sunny days in July 2012. For 200 of those clones, a second biological replicate was also sampled. Typically, leaves (leaf plastocron index 9 plus or minus 1) on a south facing branch from the upper canopy of each tree were quickly collected, wiped with a wet tissue to clean both surfaces and the leaf then fast frozen under dry ice. Leaves were kept on dry ice and shipped back to the lab and stored at −80°C until processed for analyses. Metabolites from leaf samples were lyophilized and then ground in a micro-Wiley mill (1 mm mesh size). Approximately 25 mg of each sample was twice extracted in 2.5 mL 80% ethanol (aqueous) for 24 hr with the extracts combined, and 0.5 ml dried in a helium stream. Sorbitol (75 μl of a 1 mg/mL aqueous solution) was added before extraction as an internal standard to correct for differences in extraction efficiency, subsequent differences in derivatization efficiency and changes in sample volume during heating. Metabolites in the dried sample extracts were converted to their trimethylsilyl (TMS) derivatives, and analyzed by gas chromatography-mass spectrometry, as described previously (Tschaplinski et al., 2012; Li et al., 2012). Briefly, dried extracts of metabolites were dissolved in acetonitrile followed by the addition of N-methyl-N-trimethylsilyltrifluoroacetamide (MSTFA) with 1% trimethylchlorosilane (TMCS), and samples then heated for 1 h at 70°C to generate TMS derivatives. After 2 days, aliquots were injected into an Agilent 5975C inert XL gas chromatograph-mass spectrometer (GCMS). The standard quadrupole GCMS is operated in the electron impact (70 eV) ionization mode, targeting 2.5 full-spectrum (50-650 Da) scans per second, as described previously (Tschaplinski et al., 2012). Metabolite peaks were extracted using a key selected ion, characteristic m/z fragment, rather than the total ion chromatogram, to minimize integrating co-eluting metabolites. The peak areas were normalized to the amount of internal standard (sorbitol) injected and the amount of sample extracted. A large user-created database (> 2400 spectra) of mass spectral electron impact ionization (EI) fragmentation patterns of TMS-derivatized metabolites, as well as the Wiley Registry 10th Edition combined with NIST 2014 mass spectral library, are used to identify the metabolites of interest to be quantified.

#### 3.2.2 pyMBMS Data

A commercially available molecular beam mass spectrometer (MBMS) designed specifically for biomass analysis was used for pyrolysis vapor analysis (Evans and Milne, 1987; Sykes et al., 2009; Tuskan et al., 1999). Approximately 4 mg of air dried 20 mesh biomass was introduced into the quartz pyrolysis reactor via 80 uL deactivated stainless steel Eco-Cups provided with the autosampler. Mass spectral data from m/z 30-450 were acquired on a Merlin Automation data system version 3.0 using 17 eV electron impact ionization.

The pyMBMS mz peaks were annotated as described in (Sykes et al., 2009), as done previously in (Muchero et al., 2015).

#### 3.2.3 Single Nucleotide Polymorphism Data

A dataset consisting of 28,342,758 SNPs called across 882 *P. trichocarpa* (Tuskan et al., 2006) genotypes was obtained from http://bioenergycenter.org/besc/gwas/. This dataset is derived from whole genome sequencing of undomesticated *P. trichocarpa* genotypes collected from the U.S. and Canada, and clonally replicated in common gardens (Tuskan et al., 2011). Genotypes from this population have previously been used for population genomics (Evans et al., 2014) and GWAS studies in *P. trichocarpa* (McKown et al., 2014) as well as for investigating linkage disequilibrium in the population (Slavov et al., 2012).

Whole genome resequencing was carried out on a sample 882 P. trichocarpa natural individuals to an expected median coverage of 15x using Illumina Genome Analyzer, HiSeq 2000, and HiSeq 2500 sequencing platforms at the DOE Joint Genome Institute. Alignments to the *P. trichocarpa* Nisqually-1 v.3.0 reference genome were performed using BWA v0.5.9-r16 with default parameters, followed by post-processing with the picard FixMateInformation and MarkDuplicates tools. Genetic variants were called by means of the Genome Analysis Toolkit v. 3.5.0 (GATK; Broad Institute, Cambridge, MA, USA) (McKenna et al., 2010; Van der Auwera et al., 2013). Briefly, variants were called independently for each individual using the concatenation of RealignerTargetCreator, IndelRealigner and HaplotypeCaller tools, and the whole population was combined using GenotypeGVCFs, obtaining a dataset with all the variants detected across the sample population. Biallelic SNPs were extracted using the SelectVariants tool and quality-filtered using the GATKâĂŹs machine-learning implementation Variant Quality Score Recalibration (VQSR). To this end, the tool VariantRecalibrator was used to create the recalibration file and the sensitivity tranches file. As a “truth” dataset, we used SNP calls from a population of seven female and seven male *P. trichocarpa* that had been crossed in a half diallel design. “True” SNPs were identified by the virtual absence of segregation distortion and Mendelian violations in the progeny of these 49 crosses (ca. 500 offspring in total). As a “non-true” dataset, we used the SNP calls of seven open-pollinated crosses from these 7 females (n = 90), filtered using hard-filtering methods recommended in the GATK documentation (tool: VariantFiltration; quality thresholds: QD < 1.5, FS > 75.0, MQ < 35.0, missing alleles < 0.5 and MAF > 0.05). The prior likelihoods for the true and non-true datasets were Q = 15 and Q = 10, respectively, and the variant quality annotations to define the variant recalibration space were DP, QD, MQ, MQRankSum, ReadPosRankSum, FS, SOR and InbreedingCoeff. Finally, we used the ApplyRecalibration tool on the full GWAS dataset to assign SNPs to tranches representing different levels of confidence. We selected SNPs in the tranche with true sensitivity < 90, which minimizes false positives, but at an expected cost of 10% false negatives. The final filtered dataset had a transition/transversion ratio of 2.07, compared to 1.88 for the unfiltered SNPs. To further validate the quality of these SNP calls, we compared them to an Illumina Infinium BeadArray that had been generated from a subset of this population dataset (Geraldes et al., 2013). The average match rate was 96% (±2% SD) for 641 individuals across 20,723 loci.

SNPs in this dataset were divided into different Tranches, indicating the percentage of “true” SNPs recovered. For further analysis in this study, we made use of the PASS SNPs, corresponding to the most stringent Tranche, recovering 90% of the true SNPs [ see http://gatkforums.broadinstitute.org/gatk/discussion/39/variant-quality-score-recalibration-vqsr].

VCFtools (Danecek et al., 2011) was used to extract the desired Tranche of SNPs from the VCF file and reformat it into.tfam and.tped files.

#### 3.2.4 GWAS Analysis

The metabolomics and pyMBMS data was used as phenotypes in a genome wide association analysis. The respective phenotype measured over all the genotypes were analyzed to account for potential outliers. A median absolute deviation (MAD) from the median (Leys et al., 2013) cutoff was applied to determine if a particular measurement of a given phenotype was an outlier with respect to all measurements of that phenotype across the population. To account for asymmetry, the deviation values were estimated separately for values below and above the median, respectively. The distribution of the measured values together with the distribution of their estimated deviation was analyzed and a cutoff of 5 was determined to identify putative outlier values. Phenotypes that had non-outlier measurements in at least 20 percent of the population were retained for further analysis, this was to ensure sufficient signal for the genome wide association model. This resulted in 1262 pyMBMS derived phenotypes and 818 metabolomics derived phenotypes.

To estimate the statistical significant associations between the respective phenotypes and the SNPs called across the population, we applied a linear mixed model using EMMAX Kang et al. (2010). Taking into account population structure estimated from a kinship matrix, we tested each of the respective 2080 phenotypes against the high-confidence SNPs and corrected for multiple hypotheses bias using the Benjamini-Hochberg control for false-discovery rate of 0.1 Benjamini and Hochberg (1995). This was done in parallel with a python wrapper that utilized the schwimmbad python package (Price-Whelan and Foreman-Mackey, 2017).

SNP-Phenotype GWAS networks were then pruned to only include SNPs that resided within genes, and SNPs were mapped to their respective genes, resulting in a gene-phenotype network. SNPs were determined to be within genes using the gene boundaries defined in the `Ptrichocarpa_210_v3.0.gene.gff3` from the *P. trichocarpa* version 3.0 genome assembly on Phytozome (Goodstein et al., 2012).

### 3.3. Co-Expression Network Construction

*Populus trichocarpa* (Nisqually-1) RNA-seq dataset from JGI Plant Gene Atlas project (Sreedasyam et al., unpublished) was obtained from Phytozome. This dataset consists of samples for standard tissues (leaf, stem, root and bud tissue) and libraries generated from nitrogen source study. List of sample descriptions was accessed from: https://phytozome.jgi.doe.gov/phytomine/aspect.do?name=Expression.

#### 3.3.1 Plant growth and treatment conditions

*Populus trichocarpa* (Nisqually-1) cuttings were potted in 4″ X 4″ X 5″ containers containing 1:1 mix of peat and perlite. Plants were grown under 16-h-light/8-h-dark conditions, maintained at 20-23 °C and an average of 235 *μ*mol m^−2^s^−1^ to generate tissue for (1) standard tissues and (2) nitrogen source study. Plants for standard tissue experiment were watered with McCownâĂŹs woody plant nutrient solution and plants for nitrogen experiment were supplemented with either 10mM KNO3 (NO3-plants) or 10mM NH4Cl (NH4+ plants) or 10 mM urea (urea plants). Once plants reached leaf plastochron index 15 (LPI-15), leaf, stem, root and bud tissues were harvested and immediately flash frozen in liquid nitrogen and stored at -80°C until further processing was done. Every harvest involved at least three independent biological replicates for each condition and a biological replicate consisted of tissue pooled from 3 plants.

#### 3.3.2 RNA extraction and sequencing

Tissue was ground under liquid nitrogen and high quality RNA was extracted using standard Trizol-reagent based extraction (Li and Trick, 2005). The integrity and concentration of the RNA preparations were checked initially using Nano-Drop ND-1000 (Nano-Drop Technologies) and then by BioAnalyzer (Agilent Technologies). Plate-based RNA sample prep was performed on the PerkinElmer Sciclone NGS robotic liquid handling system using Illumina’s TruSeq Stranded mRNA HT sample prep kit utilizing poly-A selection of mRNA following the protocol outlined by Illumina in their user guide: http://support.illumina.com/sequencing/sequencing_kits/truseq_stranded_mrna_ht_sample_prep_kit.html, and with the following conditions: total RNA starting material was 1 ug per sample and 8 cycles of PCR was used for library amplification. The prepared libraries were then quantified by qPCR using the Kapa SYBR Fast Illumina Library Quantification Kit (Kapa Biosystems) and run on a Roche LightCycler 480 real-time PCR instrument. The quantified libraries were then prepared for sequencing on the Illumina HiSeq sequencing platform utilizing a TruSeq paired-end cluster kit, v4, and IlluminaâĂŹs cBot instrument to generate a clustered flowcell for sequencing. Sequencing of the flowcell was performed on the Illumina HiSeq2500 sequencer using HiSeq TruSeq SBS sequencing kits, v4, following a 2×150 indexed run recipe.

#### 3.3.3 Correlation Analysis

Gene expression atlas data for *P. trichocarpa* consisting of 63 different samples were used to construct a co-expression network. Reads were trimmed using Skewer (Jiang et al., 2014). Star (Dobin et al., 2013) was then used to align the reads to the P. *trichocarpa* reference genome (Tuskan et al., 2006) obtained from Phytozome (Goodstein et al., 2012). TPM (Transcripts Per Million) expression values (Wagner et al., 2012) were then calculated for each gene. This resulted in a gene expression matrix *E* in which rows represented genes, columns represented samples and each entry *ij* represented the expression (TPM) of gene *i* in sample *j*. The Spearman correlation coefficient was then calculated between the expression profiles of all pairs of genes (i.e. all pairs of rows of the matrix E) using the mcxarray and mcxdump programs from the MCL-edge package (Van Dongen, 2008, Van Dongen 2001) available from http://micans.org/mcl/. This was performed in parallel using Perl wrappers making use of the Parallel::MPI::Simple Perl module, (Alex Gough, http://search.cpan.org/~ajgough/Parallel-MPI-Simple-0.03/Simple.pm) using compute resources at the Oak Ridge Leadership Computing Facility (OLCF).

Supplementary Figure S1A shows the distribution of Spearman correlation values for the co-expression network. An absolute threshold of 0.85 was applied.

### 3.4 Co-Methylation Network Construction

Methylation data for *P. trichocarpa* (Vining et al., 2012) re-aligned to the version 3.0 assembly of *P. trichocarpa* was obtained from Phytozome (Goodstein et al., 2012). This data consisted of MeDIP-seq (Methyl-DNA immunoprecipitation-seq) reads from 10 different *P. trichocarpa* tissues, including bud, callus, female catkin, internode explant, leaf, male catkin, phloem, regenerated internode, root and xylem tissue.

BamTools stats (Barnett et al., 2011) was used to determine basic properties of the reads in each.bam file. Samtools (Li et al., 2009) was then used to extract only mapped reads. The number of reads which mapped to each gene feature was determined using htseq-count (Anders et al., 2014). These read counts were then converted to TPM values (Wagner et al., 2012), providing a methylation score for each gene in each tissue. The TPM value for a gene *g* in a given sample was defined as:

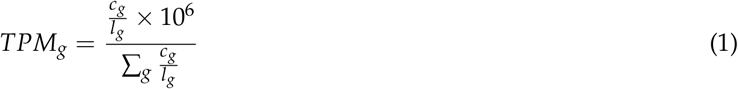

where *c*_*g*_ is the number of reads mapped to gene *g* and *l*_*g*_ is the length of gene *g* in kb, calculated by subtracting the gene start position from the gene end position, and dividing the resulting difference by 1,000. A methylation matrix *M* was then formed, in which rows represented genes, columns represented tissues and each entry *ij* represented the methylation score (TPM) of gene *i* in tissue *j*. A co-methylation network (see references (Busch et al., 2016; Akulenko and Helms, 2013; Davies et al., 2012)) was then constructed by calculating the Spearman correlation coefficient between the methylation profiles of all pairs of genes using mcxarray and mcxdump programs from the MCL-edge package (Van Dongen, 2008, 2001) http://micans.org/mcl/. Supplementary Figure S1B shows the distribution of Spearman Correlation values. An absolute threshold of 0.95 was applied.

Read counting using htseq-count, as well as Spearman correlation calculations were performed in parallel using Perl wrappers making use of the Parallel::MPI::Simple Perl module, developed by Alex Gough and available on The Comprehensive Perl Archive Network (CPAN) at http://www.cpan.org and used compute resources at the Oak Ridge Leadership Computing Facility (OLCF).

### 3.5 SNP Correlation Network Construction

The Custom Correlation Coefficient (CCC) (Climer et al., 2014b, Climer 2014a) was used to calculate the correlation between the occurrence of pairs of SNPs across the 882 genotypes. The CCC between allele *x* at position *i* and allele *y* and position *j* is defined as:

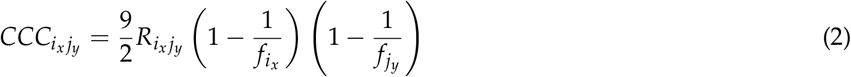

where *R*_*i*_*x*_*j*_*y*__ is the relative co-occurrence of allele *x* at position *i* and allele *y* at position *j*, *f*_*i*_*x*__ is the frequency of allele *x* at position *i* and *f*_*j*_*y*__ is the frequency of allele *y* at position *j*.

This was performed in a parallel fashion using similar computational approaches as described for the co-expression network above. The set of ~10 million SNPs was divided into 20 different blocks, and the CCC was calculated for each within-block and cross-block SNPs in separate jobs, to a total of 210 MPI jobs (Figure 2). A threshold of 0.7 was then applied. The resulting SNP correlation network was pruned to only include SNPs that resided within genes. Gene boundaries used were defined in the `Ptrichocarpa_210_v3.0.gene.gff3` from the *P. trichocarpa* version 3.0 genome assembly on Phytozome (Goodstein et al., 2012). A local LD filter was then set, retaining correlations between SNPs greater than 10kb apart. The distribution of CCC values can be seen in Supplementary Figure S1C (Supplementary Note 1).

**Figure 2:**
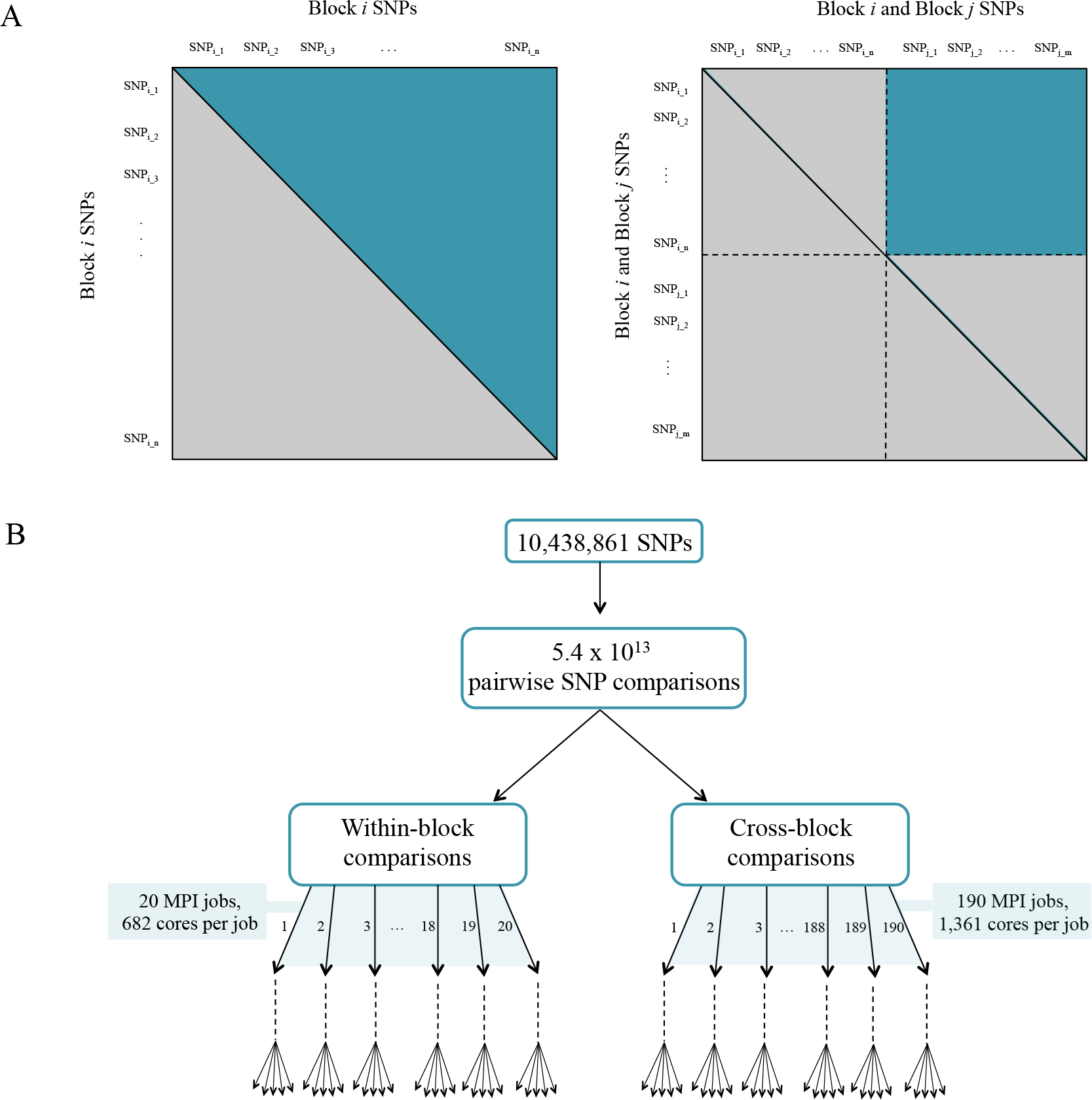
(A) Parallelization strategy for ccc calculation between all fairs of SNPs. (B) MPI jobs for within and cross-block comparisons.

### 3.6 Gene Annotation

*P. trichocarpa* gene annotations in the `Ptrichocarpa_210_v3.0.annotation_info.txt` file from the version 3.0 genome assembly were used, available on Phytozome (Goodstein et al., 2012). This included *Arabidopsis* best hits and corresponding gene descriptions, as well as GO terms (Gene Ontology Consortium, 2017; Ashburner et al., 2000) and Pfam domains (Finn et al., 2016). Genes were also assigned MapMan annotations using the Mercator tool (Lohse et al., 2014).

### 3.7 Scoring Lines of Evidence (LOE)

A scoring system was developed in order to quantify the Lines Of Evidence (LOE) linking each gene to lignin-related genes/phenotypes. The LOE scores quantify the number of lines linking each gene to lignin-related genes and phenotypes across the different network data layers. The process of defining and calculating LOE scores is described below.

#### 3.7.1 Selection of Lignin-related Genes

Lignin building blocks (monolignols) are derived from phenylalanine in the phenylpropanoid and monolignol pathways, and phenylalanine itself is produced from the shikimate pathway (Vanholme et al., 2010). To compile a list of *P. trichocarpa* genes which are related to the biosynthesis of lignin, *P. trichocarpa* genes were assigned MapMan annotations using the Mercator tool (Lohse et al., 2014). Genes in the Shikimate (MapMan bins 13.1.6.1, 13.1.6.3 and 13.1.6.4), Phenylpropanoid (MapMan bin 16.2) and Lignin/Lignan (MapMan bin 16.2.1) pathways were then selected. A list of these lignin-related genes and their MapMan annotations can be seen in Supplementary Table S1.

#### 3.7.2 Selection of Lignin-related Phenotypes

Lignin-related pyMBMS peaks, as described in Sykes et al. (2009), Davis et al. (2006) and Muchero et al. (2015) were identified among the pyMBMS GWAS hits, and are shown in Supplementary Table S2. Lignin-related metabolites and metabolites in the lignin pathway were also identified among the metabolomics GWAS hits, a list of which can be seen in Supplementary Table S3. For partially identified metabolites, additional RT and mz information can be seen in Supplementary Table S3.

#### 3.7.3 Extraction of Lignin-Related Subnetworks

Let *L*_*G*_, *L*_*M*_ and *L*_*P*_ represent our sets of lignin-related genes, metabolites and pyMBMS peaks, respectively (Supplementary Tables S1, S2 and S3). A network can be defined as *N* = (*V*, *E*) where *V* is the set of nodes and *E* is the set of edges connecting nodes in *V*. In particular, let the co-expression network be represented by *N*_*coex*_ = (*V*_*coex*_, *E*_*coex*_), the co-methylation network by *N*_*cometh*_ = (*V*_*cometh*_, *E*_*cometh*_) and the SNP correlation network by *N*_*sn p*_ = (*V*_*snp*_, *E*_*snp*_). The GWAS networks can be represented as bipartite networks *N* = (*U*, *V*, *E*) where *U* is the set of phenotype nodes, *V* is the set of gene nodes, and *E* is the set of edges, with each edge *e*_*ij*_ connecting node *i* ∈ *U* with node *j* ∈ *V*. Let the metabolomics GWAS network be represented by *N*_*metab*_ = (*U*_*metab*_, *V*_*metab*_, *E*_*metab*_) and the pyMBMS GWAS network by *N*_*pymbms*_ = (*U*_*pymbms*_, *V*_*pymbms*_, *E*_*pymbms*_). We construct the *guilt by association* subnetworks of genes connected to lignin-related genes/phenotypes as follows:

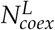 is the subnetwork of *N*_*coex*_ including the lignin related genes *l* ∈ *L*_*G*_ and their direct neighbors:

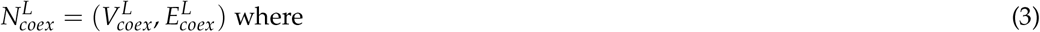

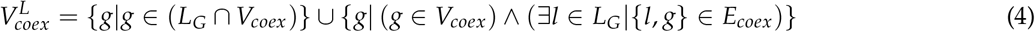

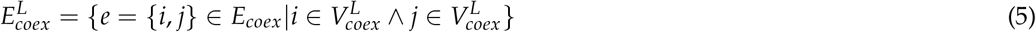
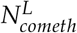 is the subnetwork of *N*_*cometh*_ including the lignin related genes *l* ∈ *L*_*G*_ and their direct neighbors:

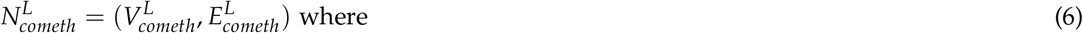

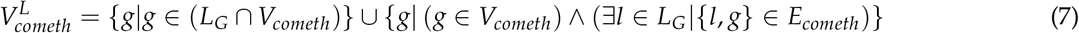

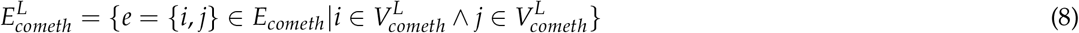
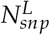 is the subnetwork of *N*_*snp*_ including the lignin related genes *l* ∈ *L*_*G*_ and their direct neighbors:

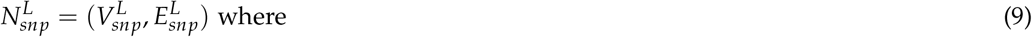

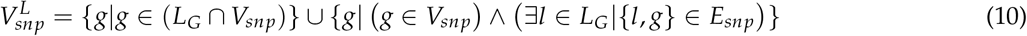

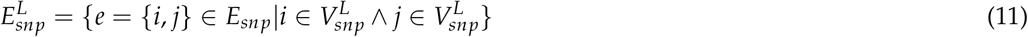
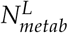 is the subnetwork of *N*_*metab*_ including the lignin related metabolites *m* ∈ *L*_*M*_ and their direct neighboring genes:

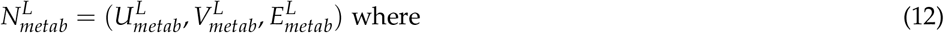

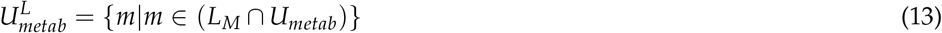

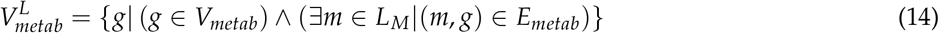

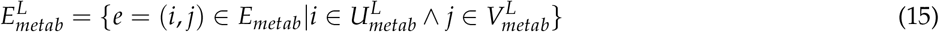
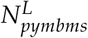 is the subnetwork of *N*_*pymbms*_ including the lignin related pyMBMS peaks *p* ∈ *L*_*P*_ and their direct neighboring genes:

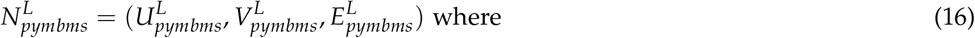

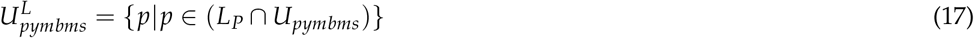

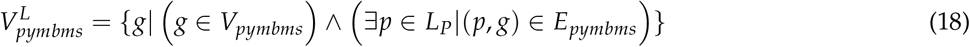

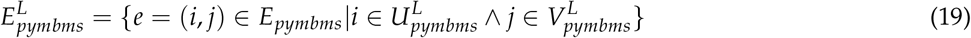

#### 3.7.4 Calculating LOE Scores

For a given gene *g*, the *degree* of that gene *D*(*g*) indicates the number of connections that the gene has in a given network. Let *D*_*coex*_(*g*), *D*_*cometh*_(*g*), *D*_*snp*_(*g*), *D*_*metab*_(*g*), *D*_*pymbms*_(*g*) represent the degrees of gene *g* in the lignin subnetworks 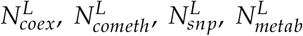 and 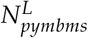, respectively. The LOE *breadth* score *LOE*_*breadth*_(*g*) is then defined as

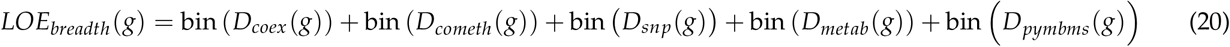

where

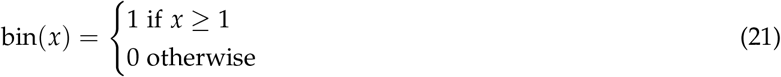

The *LOE*_*breadth*_(*g*) score indicates the number of different types of lines of evidence that exist linking gene *g* to lignin-related genes/phenotypes.

The LOE *depth* score *LOE*_*depth*_(*g*) represents the total number of lines of evidence exist linking gene *g* to lignin-related genes/phenotypes, and is defined as

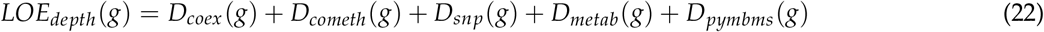

The GWAS LOE score *LOE*_*gwas*_(*g*) indicates the number of lignin-related phenotypes (metabolomic or pyMBMS) that a gene is connected to, and is defined as:

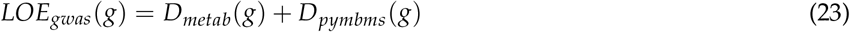

Distributions of the LOE scores can be seen in Supplementary Figure S2. Cytoscape version 3.4.0 (Shannon et al., 2003) was used for network visualization.

### 3.8 Packages Used

Networks were visualized using Cytoscape version 3.4.0 (Shannon et al., 2003). Expression, methylation, SNP correlation and GWAS diagrams were created using R (R Core Team, 2017) and various R libraries (de Vries and Ripley, 2016; Auguie, 2017; Wickham, 2007; Arnold, 2017; Wickham, 2009). Data parsing, wrappers and LOE score calculation was performed using Perl. Diagrams were edited to overlay certain text using Microsoft PowerPoint.

## 4. RESULTS AND DISCUSSION

### 4.1 Layered Networks, LOE Scores and New Potential Targets

This study involved the construction of a set of networks providing different layers of information about the relationships between genes, and between genes and phenotypes, and the development of a Lines Of Evidence scoring system (LOE scores) which integrate the information in the different network layers and quantify the number of lines of evidence connecting genes to lignin-related genes/phenotypes. The GWAS network layers provide information as to which genes are potentially involved in certain functions because they contain genomic variants significantly associated with measured phenotypes. The co-methylation and co-expression networks provide information on different layers of regulatory mechanisms within the cell. The SNP correlation network provides information about possible co-evolution relationships between genes, through correlated variants across a population.

Marking known genes and phenotypes involved in lignin biosynthesis in these networks allowed for the calculation of a set of LOE (Lines Of Evidence) scores for each gene, indicating the strength of the evidence linking each gene to lignin-related functions. The breadth LOE score indicates the number of types of lines of evidence (number of layers) which connect the gene to lignin-related genes/phenotypes, whereas the depth LOE score indicates the total number of lignin-related genes/phenotypes the gene is associated with. Individual layer LOE scores (e.g. co-expression LOE score or GWAS LOE score) indicate the number of lignin-related associations the gene has within that layer.

To select the top set of potential new candidate genes involved in lignin biosynthesis, genes which showed a number of different lines of evidence connecting them to lignin-related functions were identified by selecting genes with a LOE breadth score >= 3. Since the GWAS networks provide the highest resolution, most direct connections to lignin-related functions, it was also required that our potential new targets had a GWAS score >= 1. This provides a set of 375 new candidate genes potentially involved in lignin biosynthesis, identified through multiple lines of evidence (Supplementary Table S4). This set of Potential New Target genes will be referred to as set of PNTs. A selection of these potential new candidates below and their annotations, derived from their *Arabidopsis* best hits, will be discussed below.

### 4.2. Agamous-like Genes

Genes in the AGAMOUS-LIKE gene family are MADS-box transcription factors, many of which which have been found to play important roles in floral development (Yoo et al., 2006; Fernandez et al., 2014; Yu et al., 2017, 2004, 2002; Lee et al., 2000). Three potential AGAMOUS-LIKE (AGL) genes are found in the set of PNTs, in particular, a homolog of *Arabidopsis* AGL8 (AT5G60910, also known as FRUITFUL), a homolog of *Arabidopsis* AGL12 (AT1G71692), and a homolog of *Arabidopsis* AGL24 (AT4G24540) and AGL22 (AT2G22540).

The first potential AGL gene in our set of PNTs is Potri.012G062300, with a breadth score of 3 and a GWAS score of 2 (Figure 3A), whose best *Arabidopsis thaliana* hit is AGL8 (AT5G60910). It has GWAS associations with a lignin-related metabolite (quinic acid) and a lignin pyMBMS peak (syringol) (Figure 3C, Table 1) and is co-methylated with three lignin-related genes (Figure 3B, Table 3). There is thus strong evidence for the involvement of *P. trichocarpa* AGL8 in the regulation of lignin-related functions. There is literature evidence that supports the hypothesis of AGL8’s involvement in the regulation of lignin biosynthesis. A patent exists for the use of AGL8 expression in reducing the lignin content of plants (Yanofsky et al., 2004). The role of AGL8 (FUL) was described in Ferrándiz et al. (2000), in which they investigated the differences in lignin deposition in transgenic plants in which AGL8 is constitutively expressed, loss-of-function AGL8 mutants and wild-type *Arabidopsis* plants (Ferrándiz et al., 2000). In wild-type plants, a single layer of valve cells were lignified. In loss-of-function AGL8 mutants, all valve mesophyl cell layers were lignified, while in the transgenic plants, constitutive expression of AGL8 resulted in loss of lignified cells (Ferrándiz et al., 2000). This study thus showed the involvement of AGL8 in fruit lignification during fruit development.

**Figure 3:**
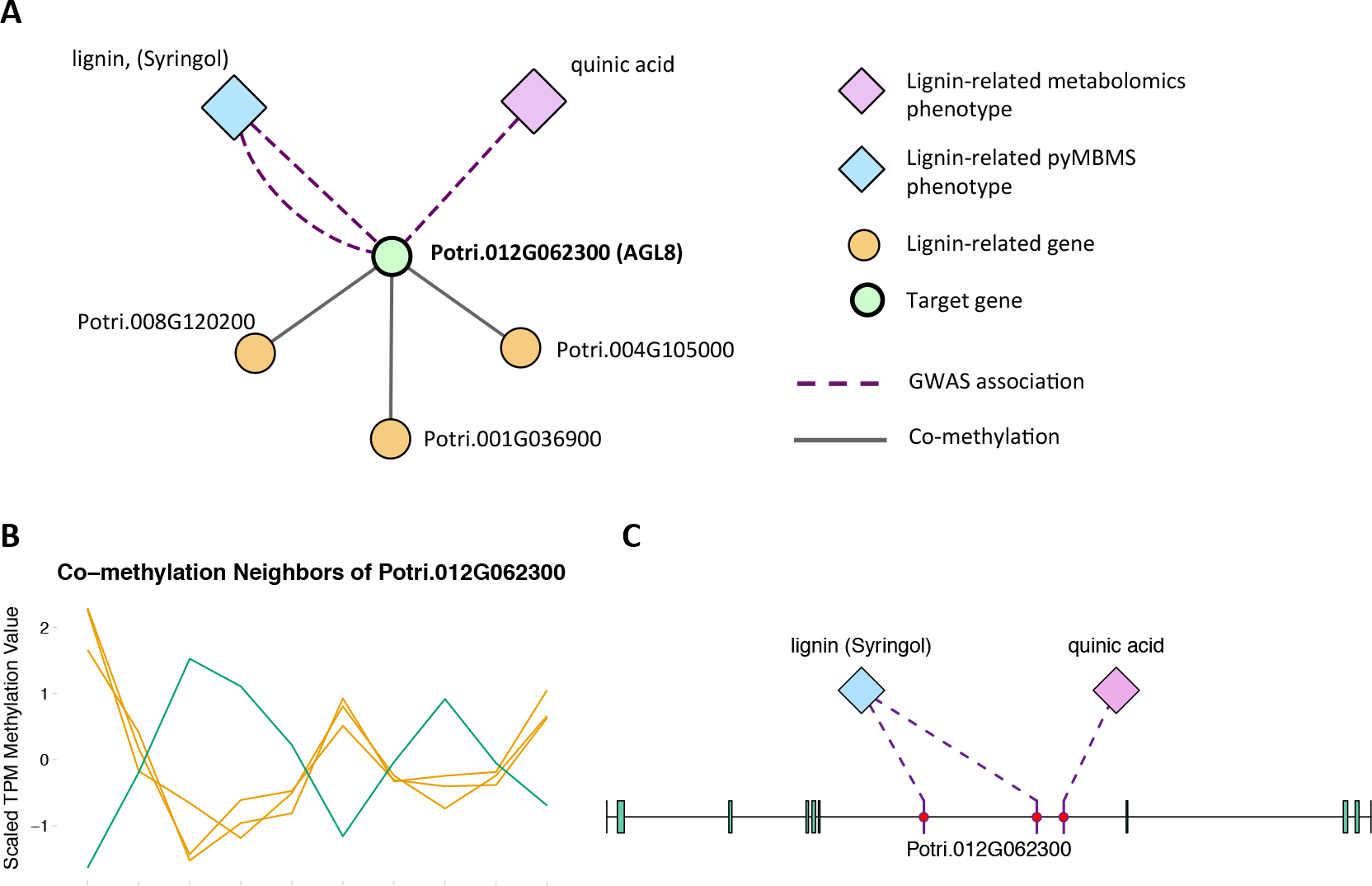
(A) Lines of Evidence for Potri.012G062300 (homolog of Arabidopsis AGL8). (B) Co-methylation of Potri.012G062300 with three lignin-related genes (Table 3) The green line represents potential target Potri.012G062300 and yellow lines represent lignin-related genes. (C) GWAS associations of Potri.012G062300 with a lignin-related metabolite and a lignin-related pyMBMS peak (Table 1).

**Table 1:** GWAS associations for select new potential target genes, indicating the SNP(s) within the potential new target gene which are associated with the lignin-related phenotype(s). Additional RT and mz information for partially identified metabolites can be seen in Supplementary Table S3.

| Source SNP | Source Gene | Target Phenotype |
| --- | --- | --- |
| **GWAS Associations for Potri.012G062300 (AGL8, AT5G60910)** |  |  |
| 12:6952245 | Potri.012G062300 | quinic acid |
| 12:6948543 | Potri.012G062300 | lignin (Syringol) |
| 12:6951532 | Potri.012G062300 | lignin (Syringol) |
| **GWAS Associations for Potri.013G102600 (AGL12, AT1G71692)** |  |  |
| 13:11604094 | Potri.013G102600 | 3-O-caffeoyl-quinic acid |
| 13:11606331 | Potri.013G102600 | coumaroyl-tremuloidin |
| 13:11600422 | Potri.013G102600 | coumaroyl-tremuloidin |
| 13:11601236 | Potri.013G102600 | hydroxyphenyl lignan glycoside |
| **GWAS Associations for Potri.007G115100 (AGL22, AT2G22540/AGL24, AT4G24540)** |  |  |
| 07:13650194 | Potri.007G115100 | caffeoyl conjugate |
| 07:13651354 | Potri.007G115100 | caffeoyl conjugate |
| 07:13642539 | Potri.007G115100 | caffeoyl conjugate |
| 07:13639923 | Potri.007G115100 | lignin, syringyl (Syringaldehyde) |
| **GWAS Associations for Potri.009G053900 (MYB46, AT5G12870)** |  |  |
| 09:5768381 | Potri.009G053900 | hydroxyphenyl lignan glycoside |
| **GWAS Associations for Potri.010G141000 (MYB111, AT5G49330)** |  |  |
| 10:15273000 | Potri.010G141000 | benzoyl-salicylate caffeic acid conjugate |
| **GWAS Associations for Potri.006G170800 (MYB36, AT5G57620)** |  |  |
| 06:17847162 | Potri.006G170800 | mz 297, RT 17.14 |
| **GWAS Associations for Potri.016G078600 (CPSRP54, AT5G03940)** |  |  |
| 16:5995136 | Potri.016G078600 | caffeoyl conjugate |
| 16:5995136 | Potri.016G078600 | feruloyl conjugate |
| 16:5996083 | Potri.016G078600 | salicyl-coumaroyl-glucoside |
| 16:5999408 | Potri.016G078600 | salicyl-coumaroyl-glucoside |
| 16:5999474 | Potri.016G078600 | salicyl-coumaroyl-glucoside |
| 16:6000236 | Potri.016G078600 | salicyl-coumaroyl-glucoside |

There is evidence of other AGAMOUS-LIKE genes affecting lignin content. A study by Gimenez *et al*. (2010) investigated TALG1, an AGAMOUS-LIKE gene in tomato, and found that TAGL1 RNAi-silenced fruits showed increased lignin content, and increased expression levels of lignin biosynthesis genes (Giménez et al., 2010). A recent study by Cosio *et al*. (2017) showed that AGL15 in *Arabidopsis* is also involved in regulating lignin-related functions, in that AGL15 binds to the promotor of peroxidase PRX17, and regulates its expression (Cosio et al., 2017). In addition, PRX17 loss of function mutants had reduced lignin content (Cosio et al., 2017).

There is thus compelling evidence that various AGAMOUS-LIKE genes are involved in regulating lignin biosynthesis/deposition in plants. Two other AGAMOUS-like genes are seen in the set of PNTs, namely a homolog of *Arabidopsis* AGL12 (Potri.013G102600) and a homolog of *Arabidopsis* AGL22/AGL24 (Potri.007G115100). Potri.013G102600 (AGL12) has GWAS associations with three lignin-related metabolites, namely hydroxyphenyl lignan glycoside, coumaroyl-tremuloidin and 3-O-caffeoyl-quinate (Figure 4A, Figure 4B, Table 1). It is co-expressed with four lignin-related genes including two caffeoyl coenzyme A O-methyltransferases, a caffeate O-methyltransferase and a ferulic acid 5-hydroxylase (Figure 4A, Figure 4C, Table 2) and it is co-methylated with four other lignin-related genes (Figure 4A, Figure 4D, Table 3). Potri.007G115100 (AGL22/AGL24) has GWAS associations with the syringaldehyde pyMBMS phenotype and a caffeoyl conjugate metabolite (Figure 5A, Figure 5B, Table 1). It also has SNP correlations with a laccase and a nicotinamidase (Figure 5A, Figure 5C, Figure 5D, Table 4, Supplementary Table S5). The combination of the multiple lines of multi-omic evidence thus suggest the involvement of *P. trichocarpa* homologs of *A. thaliana* AGL22/AGL24 and AGL12 in regulating lignin biosynthesis.

**Figure 4:**
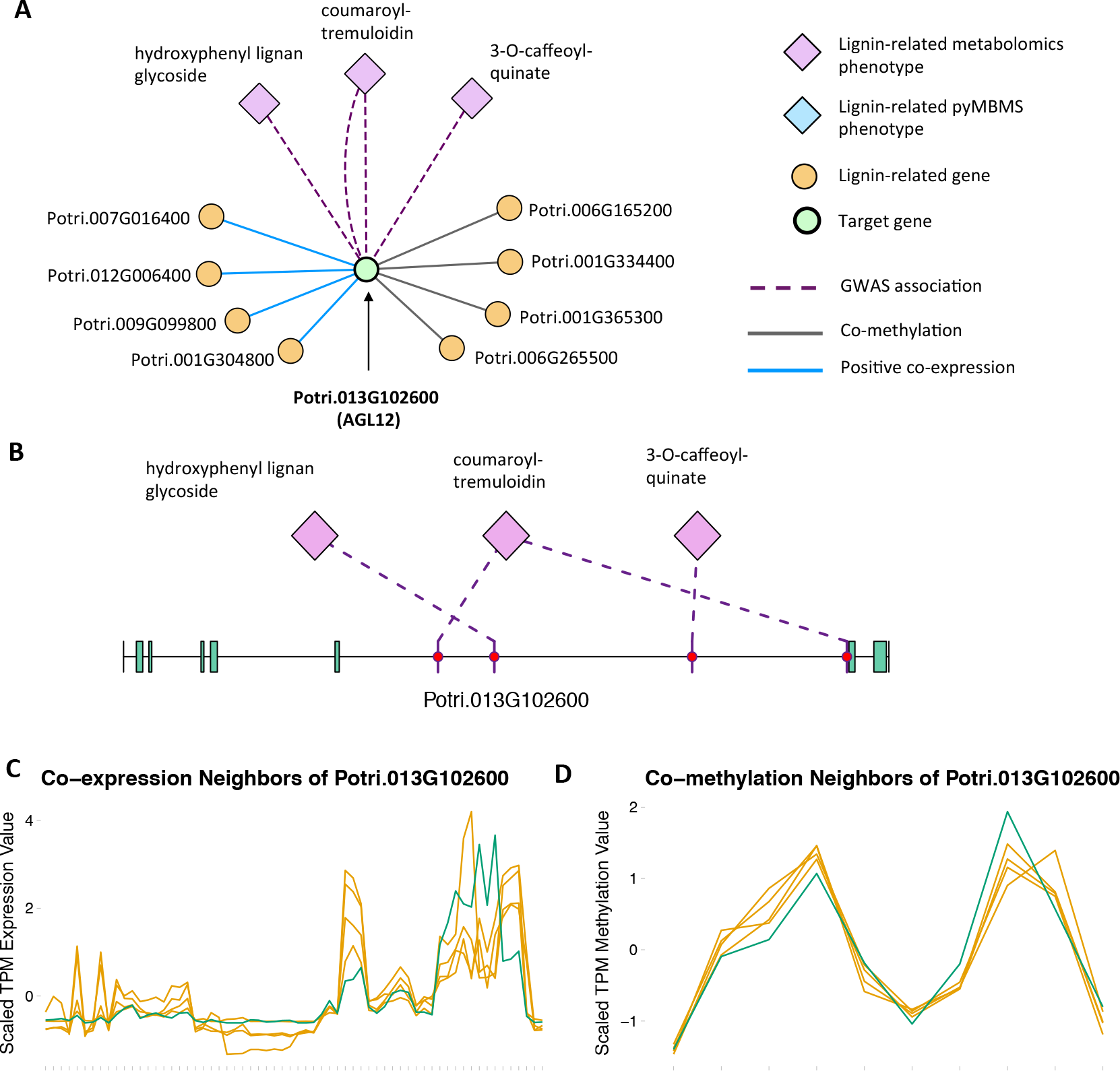
(A) Lines of Evidence for Potri.013G102600 (homolog of Arabidopsis AGL12). (B) GWAS associations of Potri.013G102600 with three lignin-related metabolites (Table 1). (C) Co-expression of Potri.013G102600 with three lignin-related genes (Table 2). (D) Co-metliylation of Potri.013G102600 with four lignin-related genes (Table 3). In line plots, the green lines represent potential target Potri.013G102600 and yellow lines represent lignin-related genes.

**Figure 5:**
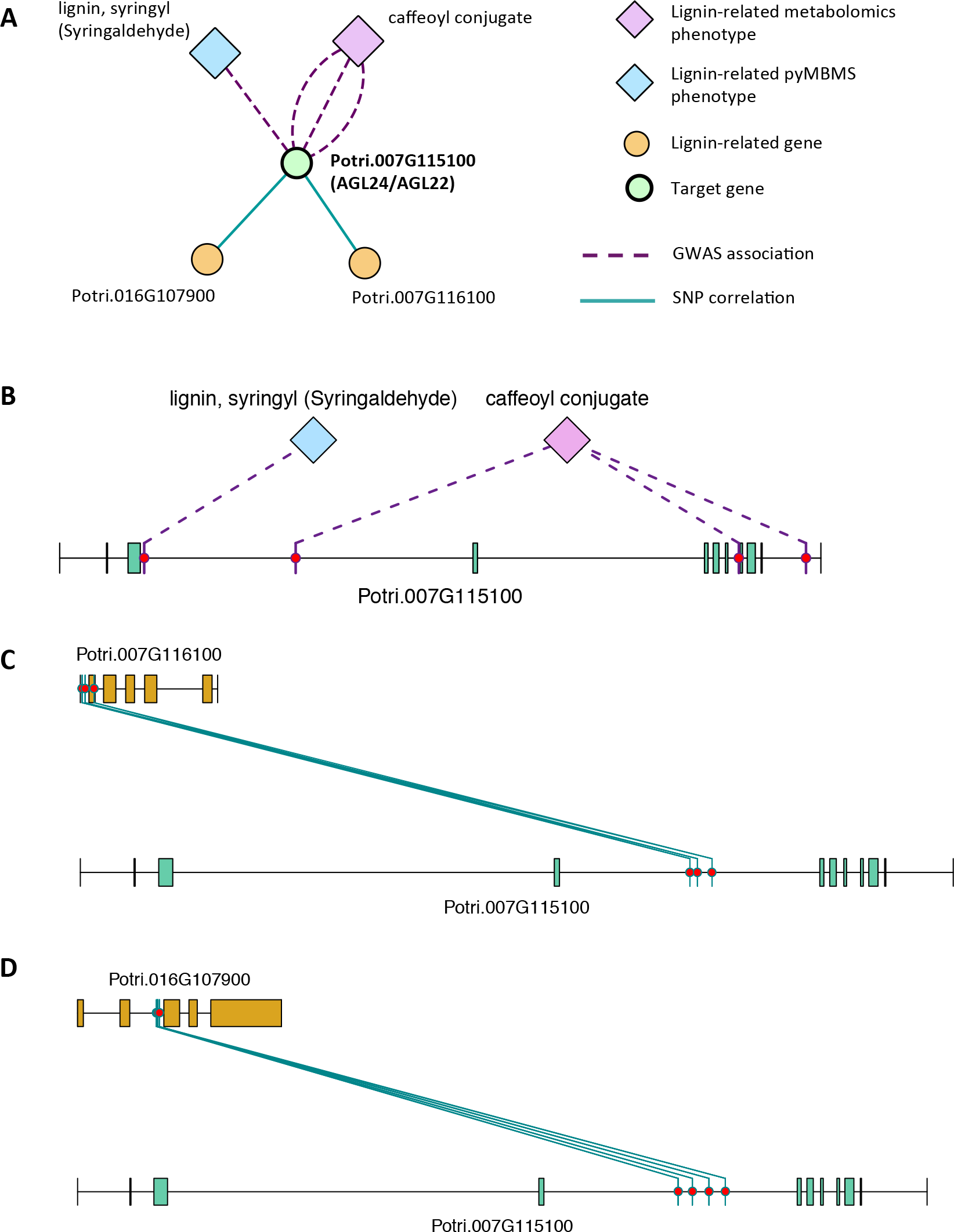
(A) Lines of Evidence for Potri.007G115100 ( homolog of Arabidopsis AGL22/24). (B) GWAS associations of Potri.007G115100 with a lignin-related metabolite and a lignin-related pyMBMS peak (Table 1). (C,D) Correlations of SNPs in Potri.007G115100 with SNPs in two lignin-related genes (Table 4, Supplementary Table S5).

**Table 2:**
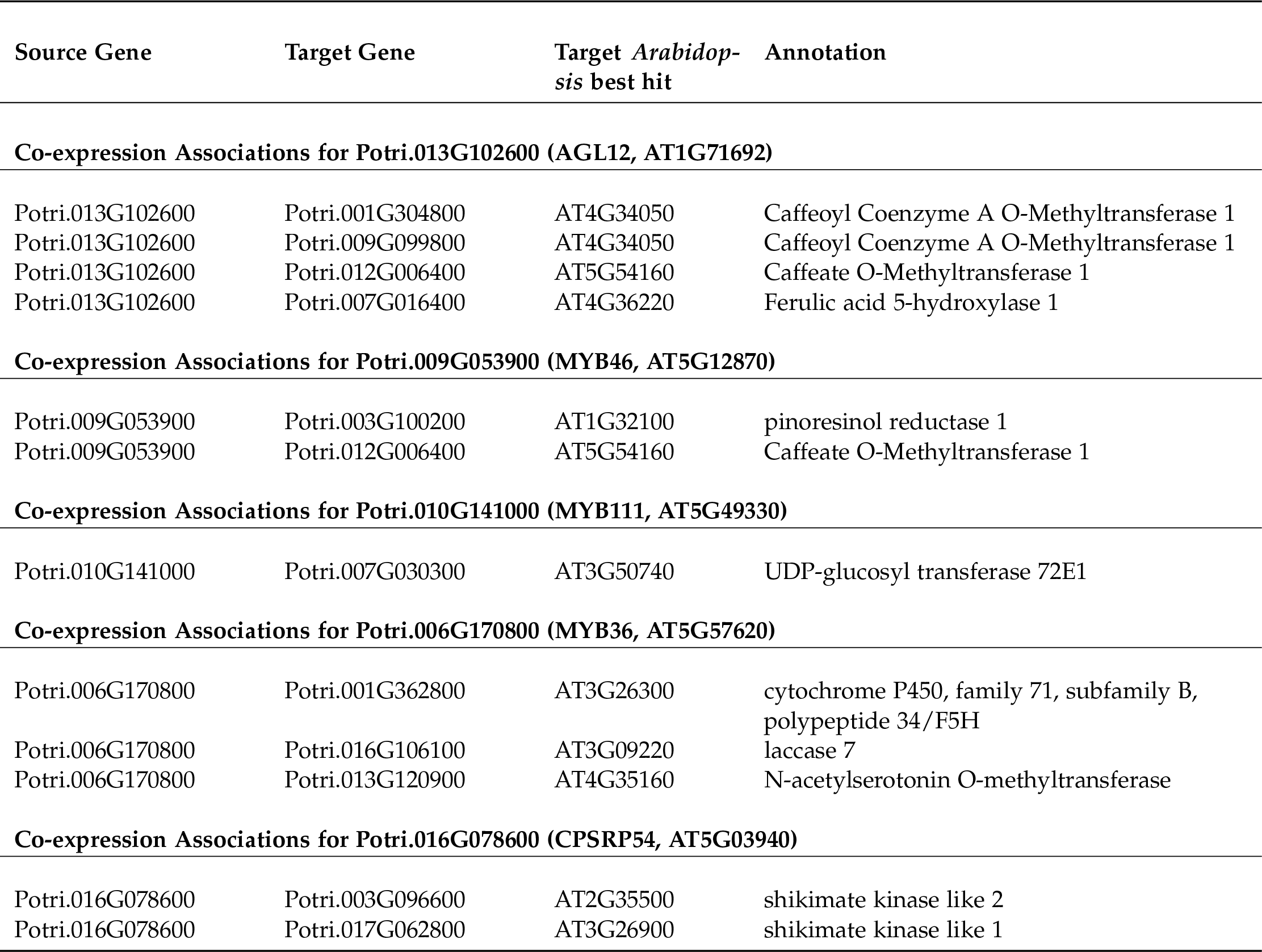
Co-expression associations for select new potential target genes. Annotations are derived from best Arabidopsis hit descriptions and GO terms and in some cases MapMan annotations.

### 4.3. MYB Transcription Factors

MYB proteins contain the conserved MYB DNA-binding domain, and usually function as transcription factors. R2R3-MYBs have been found to regulate various functions, including flavonol biosynthesis, anthocyanin biosynthesis, lignin biosynthesis, cell fate and developmental functions (Dubos et al., 2010). The set of PNTs contains several genes which are homologs of *Arabidopsis* MYB transcription factors, including homologs of *Arabidopsis* MYB66/MYB3, MYB46, MYB36 and MYB111.

There is already existing literature evidence for how some of these MYBs affect lignin biosynthesis. Liu et al. (2015) reviews the involvement of MYB transcription factors in the regulation of phenylpropanoid metabolism. MYB3 in *Arabidopsis* is known to repress phenylpropanoid biosynthesis (Zhou et al., 2017a), and a *P. trichocarpa* homolog of MYB3 is found in our set of potential new targets. Another potential new target is the *P. trichocarpa* homolog of *Arabidopsis* MYB36 (Potri.006G170800) which is connected to lignin-related functions through multiple lines of evidence (Figure 6). In *Arabidopsis*, MYB36 has been found to regulate the local deposition of lignin during casparian strip formation, and *myb36* mutants exhibit incorrectly localized lignin deposition (Kamiya et al., 2015).

**Figure 6:**
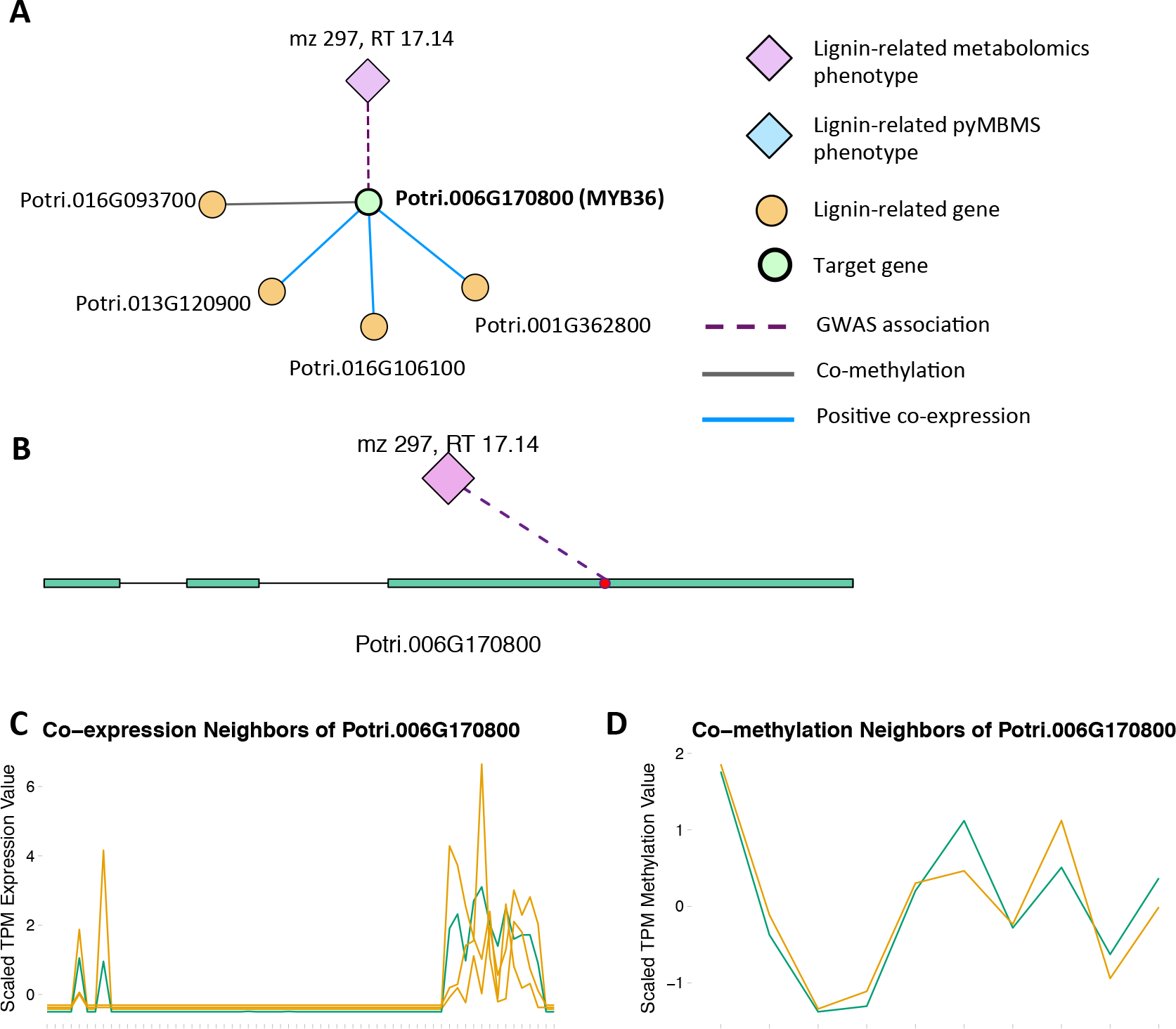
(A) Lines of Evidence for Potri.006G170800 (homolog of Arabidopsis MYB36). (B)GWAS associations of Potri.006G170800 with a lignin-related metabolite (Table 1). (C) Co-expression of Potri.006G170800 with three lignin-related genes (Table 2). (D) Co-methylation of Potri.006G170800 with a lignin-related gene (Table 3). In line plots, the green lines represent potential target Potri.006G170800 and yellow lines represent lignin-related genes.

**Table 3:** Co-methylation associations for select new potential target genes. Annotations are derived from best Arabidopsis hit descriptions and GO terms and in some cases MapMan annotations.

| Source Gene | Target Gene | Target *Arabidopsis* best hit | Annotation |
| --- | --- | --- | --- |
| **Co-methylation Associations for Potri.012G062300 (AGL8, AT5G60910)** |  |  |  |
| Potri.012G062300 | Potri.001G036900 | AT3G21240 | 4-coumarate:CoA ligase 2 |
| Potri.012G062300 | Potri.008G120200 | AT1G68540 | Cinnamoyl CoA reductase-like 6 |
| Potri.012G062300 | Potri.004G105000 | AT5G14700 | (NAD(P)-binding Rossmann-fold superfamily protein, cinnamoyl-CoA reductase activity/CCR1 |
| **Co-methylation Associations for Potri.013G102600 (AGL12, AT1G71692)** |  |  |  |
| Potri.013G102600 | Potri.001G334400 | AT5G63380 | 4-coumarate-CoA ligase activity /4CL |
| Potri.013G102600 | Potri.001G365300 | AT3G26300 | cytochrome P450, family 71, subfamily B, polypeptide 34/F5H |
| Potri.013G102600 | Potri.006G265500 | AT5G10820 | Major facilitator superfamily protein/Phenylpropanoid pathway |
| Potri.013G102600 | Potri.006G165200 | AT2G19070 | spermidine hydroxycinnamoyl transferase |
| **Co-methylation Associations for Potri.009G053900 (MYB46, AT5G12870)** |  |  |  |
| Potri.009G053900 | Potri.008G196100 | AT3G06350 | bi-functional dehydroquinate-shikimate dehydrogenase enzyme |
| Potri.009G053900 | Potri.002G018300 | AT4G39330 | cinnamyl alcohol dehydrogenase 9 |
| Potri.009G053900 | Potri.004G102000 | AT4G05160 | 4-coumarate-CoA ligase activity/4CL) |
| Potri.009G053900 | Potri.008G136600 | AT1G67980 | caffeoyl-CoA 3-O-methyltransferase |
| **Co-methylation Associations for Potri.010G141000 (MYB111, AT5G49330)** |  |  |  |
| Potri.010G141000 | Potri.008G196100 | AT3G06350 | bi-functional dehydroquinate-shikimate dehydrogenase enzyme |
| Potri.010G141000 | Potri.004G102000 | AT4G05160 | 4-coumarate-CoA ligase activity/4CL |
| Potri.010G141000 | Potri.008G074500 | AT5G34930 | arogenate dehydrogenase |
| Potri.010G141000 | Potri.005G028000 | AT5G48930 | hydroxycinnamoyl-CoA shikimate/quinic acid hydroxycinnamoyl transferase |
| Potri.010G141000 | Potri.018G100500 | AT2G23910 | NAD(P)-binding Rossmann-fold superfamily protein, cinnamoyl-CoA reductase activity/CCR1 |
| Potri.010G141000 | Potri.010G230200 | AT1G20510 | OPC-8:0 CoA ligase1, 4-coumarate-CoA ligase activity/4CL |
| **Co-methylation Associations for Potri.006G170800 (MYB36, AT5G57620)** |  |  |  |
| Potri.006G170800 | Potri.016G093700 | AT4G05160 | AMP-dependent synthetase and ligase family, 4-coumarate-CoA ligase activity/4CL |
| **Co-methylation Associations for Potri.016G078600 (CPSRP54, AT5G03940)** |  |  |  |
| Potri.016G078600 | Potri.014G135500 | AT3G06350 | bi-functional dehydroquinate-shikimate dehydrogenase enzyme |

MYB46 is known to be a regulator of secondary cell wall formation (Zhong et al., 2007). Overexpression of MYB46 in *Arabidopsis* activates lignin, cellulose and xylan biosynthesis pathways (Zhong et al., 2007). The MYB46 homolog in *P. trichocarpa*, Potri.009G053900, is connected to lignin-related functions through multiple lines of evidence (Figure 7A), including a GWAS association with a hydroxyphenyl lignan glycoside (Figure 7E, Table 1), co-expression with pinoresinol reductase 1 and caffeate O-methyltransferase 1 (Figure 7F, Table 2) and co-methylation with dehydroquinate-shikimate dehydrogenase enzyme, cinnamyl alcohol dehydrogenase 9, 4-coumarate-CoA ligase activity/4CL) and caffeoyl-CoA 3-O-methyltransferase (Figure 7G, Table 3).

**Figure 7:**
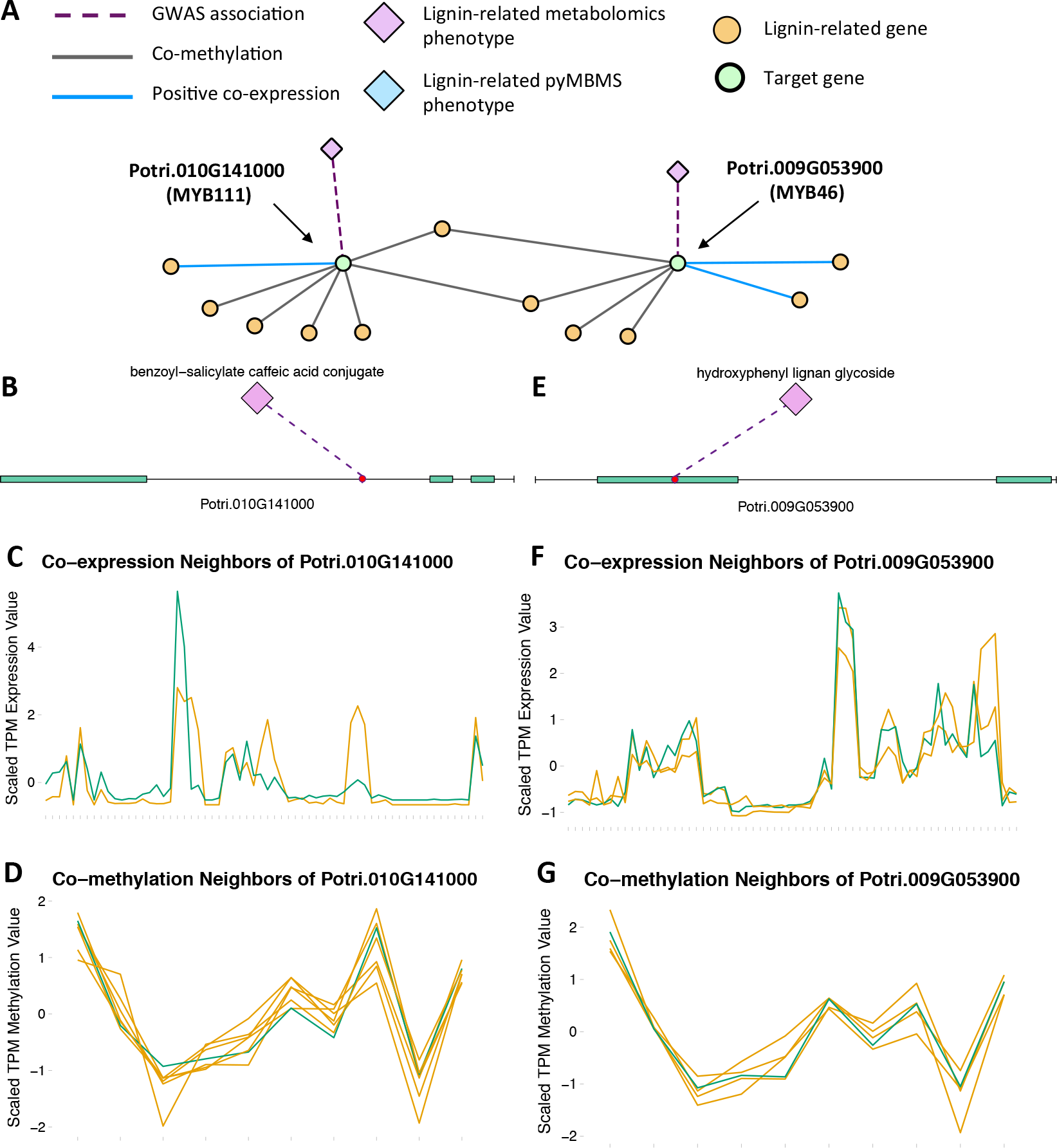
(A) Lines of Evidence for Potri.009G053900 (homolog of Arabidopsis MYB46) and Potri.010G141000 (homolog of Arabidopsis MYB111). (B) GWAS associations of Potri.010G141000 with a lignin-related metabolite (Table 1). (C) Co-expression of Potri.010G141000 with a lignin-related gene (Table 2). (D) Co-methylation of Potri.010G141000 with six lignin-related genes (Table 3). (E) GW4S associations of Potri.009G053900 with a lignin-related metabolite (Table 1). (F) Co-expression of Potri.009G053900 with two lignin-related genes (Table 2). (G) Co-methylation of Potri.009G053900 with four lignin-related genes (Table 3). In line plots, the green lines represent potential targets Potri.009G053900/Potri.010G141000 and yellow lines represent lignin-related genes.

A MYB transcription factor in the set of PNTs which has, to our knowledge, not yet been directly associated with lignin biosynthesis is MYB111 (Figure 7A-D). However, with existing literature evidence, one can hypothesize that MYB111 can alter lignin content by redirecting carbon flux from flavonoids to monolignols. There is evidence that MYB111 is involved in crosstalk between lignin and flavonoid pathways. Monolignols and flavonoids are both derived from phenylalanine through the phenylpropanoid pathway (Liu et al., 2015). There is crosstalk between the signalling pathways of ultraviolet-B (UV-B) stress and biotic stress pathways (Schenke et al., 2011). In the study by Schenke et al. (2011), it was shown that under UV-B light stress, *Arabidopsis* plants produce flavonols as a UV protectant. Also, simultaniously applying the bacterial elicitor flg22, which simulates biotic stress, repressed flavonol biosynthesis genes and induced production of defense compounds including camalexin and scopoletin, as well as lignin, which provides a physical barrier preventing pathogens’ entry (Schenke et al., 2011). This crosstalk involved regulation by MYB12 and MYB4 (Schenke et al., 2011). This study by Schenke et al. (2011) was performed using cell cultures. A second study (Zhou et al., 2017b) used *Arabidopsis* seedlings, and found that MYB111 may be involved in the crosstalk in planta (Zhou et al., 2017b). The multiple lines of evidence connecting the *P. trichocarpa* homolog of *Arabidopsis* MYB111 (Potri.010G141000) to lignin related functions, in combination with the above literature evidence suggests the involvement this gene in the regulation of lignin biosynthesis by redirecting carbon flux from flavonol biosynthesis to monolignol biosynthesis, as part of the crosstalk between UV-B protection and biotic stress signalling pathways.

### 4.4. Chloroplast Signal Recognition Particle

Potri.016G078600, a homolog of the *Arabidopsis* chloropast signal recognition particle cpSRP54 occurs in the set of PNTs (Figure 8). It has a GWAS LOE score of 3, through GWAS associations with salicyl-coumaroyl-glucoside, a caffeoyl conjugate and a feruloyl conjugate (Figure 8B, Table 1, Supplementary Table S4). It also has a breadth score of 4, indicating that it is linked to lignin-related genes/phenotypes though 4 different types of associations (Figure 8). CpSRP54 gene has been found to regulate carotenoid accumulation in *Arabidopsis* (Yu et al., 2012). CpSRP54 and cpSRP43 form a “transit complex” along with a light-harvesting chlorophyll a/b-binding protein (LHCP) family member to transport it to the thylakoid membrane (Groves et al., 2001; Schünemann, 2004). A study in *Arabidopsis* found that cpSRP43 mutants had reduced lignin content (Klenell et al., 2005). Since CpSRP54 regulates carotenoid accumulation, and cpSRP43 appears to affect lignin content, it is possible that chloroplast signal recognition particles affect lignin and carotenoid content through flux through the phenylpropanoid pathway, the common origin of both of these compounds. In fact, a gene mutation *cue1* which causes LHCP underexpression also results in reduced aromatic amino acid biosynthesis (Streatfield et al., 1999). These multiple lines of evidence, combined with the above cited literature suggests that chloroplast signal recognition particles in *P. trichocarpa* could potentially influence lignin content.

**Figure 8:**
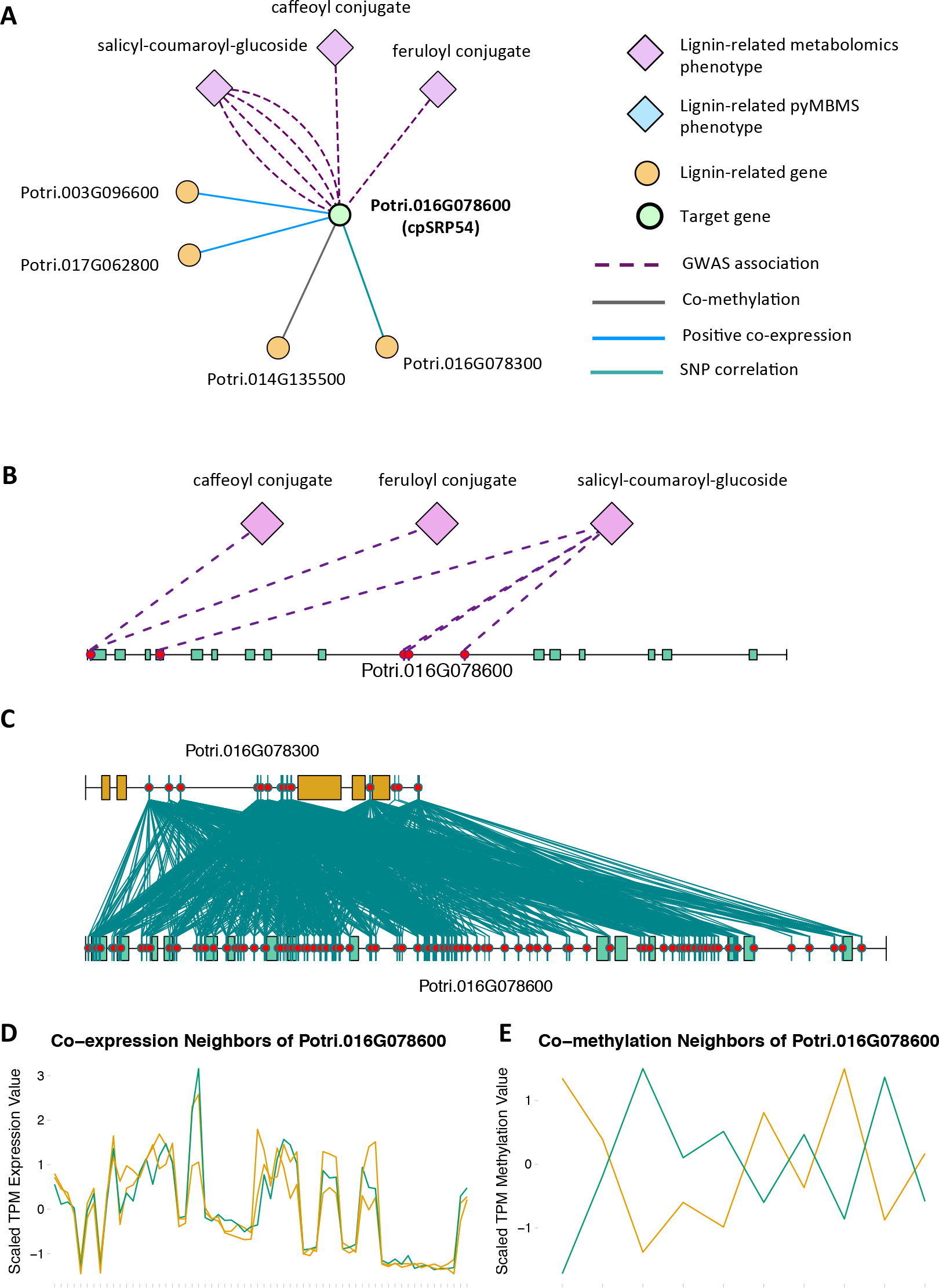
(A) Lines of Evidence for Potri.016G078600 (homolog of Arabidopsis cpSRP54). (B)GWAS associations of Potri.016G078600 with three lignin-related metabolite (Table 1). (C) Correlations of SNPs within Potri.016G078600 with SNPs in a lignin-related gene (Table 4). (D) Co-expression ofPotri.016G078600 with two lignin-related genes (Table 2). (E) Co-methylation of Potri.016G078600 with a lignin-related gene (Table 3). In line plots, the green lines represent potential target Potri.016G078600 and yellow lines represent lignin-related genes.

**Table 4:** SNP correlation associations for select new potential target genes. Annotations are derived from best Arabidopsis hit descriptions and GO terms and in some cases MapMan annotations.

| Source gene | Target gene | Target<br>best hit | *Arabidopsis* | Annotation |
| --- | --- | --- | --- | --- |
| **SNP Correlations for Potri.007G115100 (AGL22, AT2G22540/AGL24, AT4G24540)** |  |  |  |  |
| Potri.007G115100 | Potri.007G116100 | AT2G22570 |  | nicotinamidase 1 |
| Potri.007G115100 | Potri.016G107900 | AT3G09220 |  | laccase 7 |
| **SNP Correlations for Potri.016G078600 (CPSRP54, AT5G03940)** |  |  |  |  |
| Potri.016G078600 | Potri.016G078300 | AT4G37970 |  | cinnamyl alcohol dehydrogenase 6 |

### 4.5. Concluding Remarks

This study made use of high-resolution GWAS data, combined with co-expression, co-methylation and SNP correlation networks in a multi-omic, data layering approach which has allowed the identification of new potential target genes involved in lignin biosynthesis/regulation. Various literature evidence supports the involvement of many of these new target genes in lignin biosynthesis/regulation, and these are suggested for future validation for involvement in the regulation of lignin biosynthesis. The data layering technique and LOE scoring system developed can be applied to other omic data types to assist in the generation of new hypotheses surrounding various functions of interest.

## CONFLICT OF INTEREST STATEMENT

The authors declare that the research was conducted in the absence of any commercial or financial relationships that could be construed as a potential conflict of interest.

## AUTHOR CONTRIBUTIONS

DW calculated methylation TPM values, constructed the networks, developed the scoring technique, performed the data layering and scoring analysis and interpreted the results, PJ performed the outlier analysis and GWAS, MS mapped gene expression atlas reads and calculated gene expression TPM values, SD, GT and WM lead the effort on constructing the GWAS population, TJT led the leaf sample collection for GCMS-based metabolomic analyses, identified the peaks, and summarized the metabolomics data, MZM collected the leaf samples and manually extracted the metabolite data, NZ conducted leaf sample preparation, extracted and derivatized and analyzed the metabolites by GCMS, PR aided in peak extraction, JS and AS generated the gene expression atlas data, SD and DMS generated the SNP calls, RS generated the pyMBMS data, DJ conceived of and supervised the project, generated MapMan annotations and edited the manuscript, DW, PJ, SD, DMS, RS, TJT, JS and AS wrote the manuscript.

## Funding

Funding provided by The BioEnergy Science (BESC) and The Center for Bioenergy Innovation (CBI). U.S. Department of Energy Bioenergy Research Centers supported by the Office of Biological and Environmental Research in the DOE Office of Science.

This research was also supported by the Department of Energy Laboratory Directed Research and Development funding (7758), at the Oak Ridge National Laboratory. Oak Ridge National Laboratory is managed by UT-Battelle, LLC,for the US DOE under contract DE-AC05-00OR22725.

This research used resources of the Oak Ridge Leadership Computing Facility (OLCF) and the Compute and Data Environmentfor Science (CADES) at the Oak Ridge National Laboratory, which is supported by the Office of Science of the U.S. Department of Energy under Contract No. DE-AC05-00OR22725.

Support for the Poplar GWAS dataset is provided by the U.S. Department of Energy, Office of Science Biological and Environmental Research (BER) via the Bioenergy Science Center (BESC) under Contract No. DE-PS02-06ER64304. The Poplar GWAS Project used resources of the Oak Ridge Leadership Computing Facility and the Compute and Data Environment for Science at Oak Ridge National Laboratory, which is supported by the Office of Science of the U.S. Department of Energy under Contract No. DE-AC05-00OR22725.

The JGI Plant Gene Atlas project conducted by the U.S. Department of Energy Joint Genome Institute was supported by the Office of Science of the U.S. Department of Energy under Contract No. DE-AC02-05CH11231. Full Gene Atlas data sets are available at http://phytozome.jgi.doe.gov.

## ACKNOWLEDGMENTS

The authors would like to acknowledge Nancy Engle, David Weston, Ryan Aug, KC Cushman, Lee Gunter and Sara Jawdy for the metabolomics sample collection, Carissa Bleker for working on the GWAS and outlier analysis, Mark Davis for the pyMBMS data and the Department of Energy Joint Genome Institute (JGI) for sequencing.

## Supplementary Material

### 1. SUPPLEMENTARY NOTES

#### Note S1: Constructing Samples CCC Distribution

Printing out the complete result set of all possible pairwise comparisons of ~10,000,000 SNPs would require more disk space than was possibly available. In order to construct an approximate distribution of the CCC values, we selected a random subset of 100,000 SNPs and calculated the CCC correlation between all pairs of these SNPs, storing all correlation values. This sampled set of correlations was used to compute the CCC distribution. Thereafter, the CCC was calculated between all pairs of all ~10,000,000 SNPs. Only correlations meeting a threshold of 0.7 were stored.

### 2. SUPPLEMENTARY FIGURES

**Figure S1:**
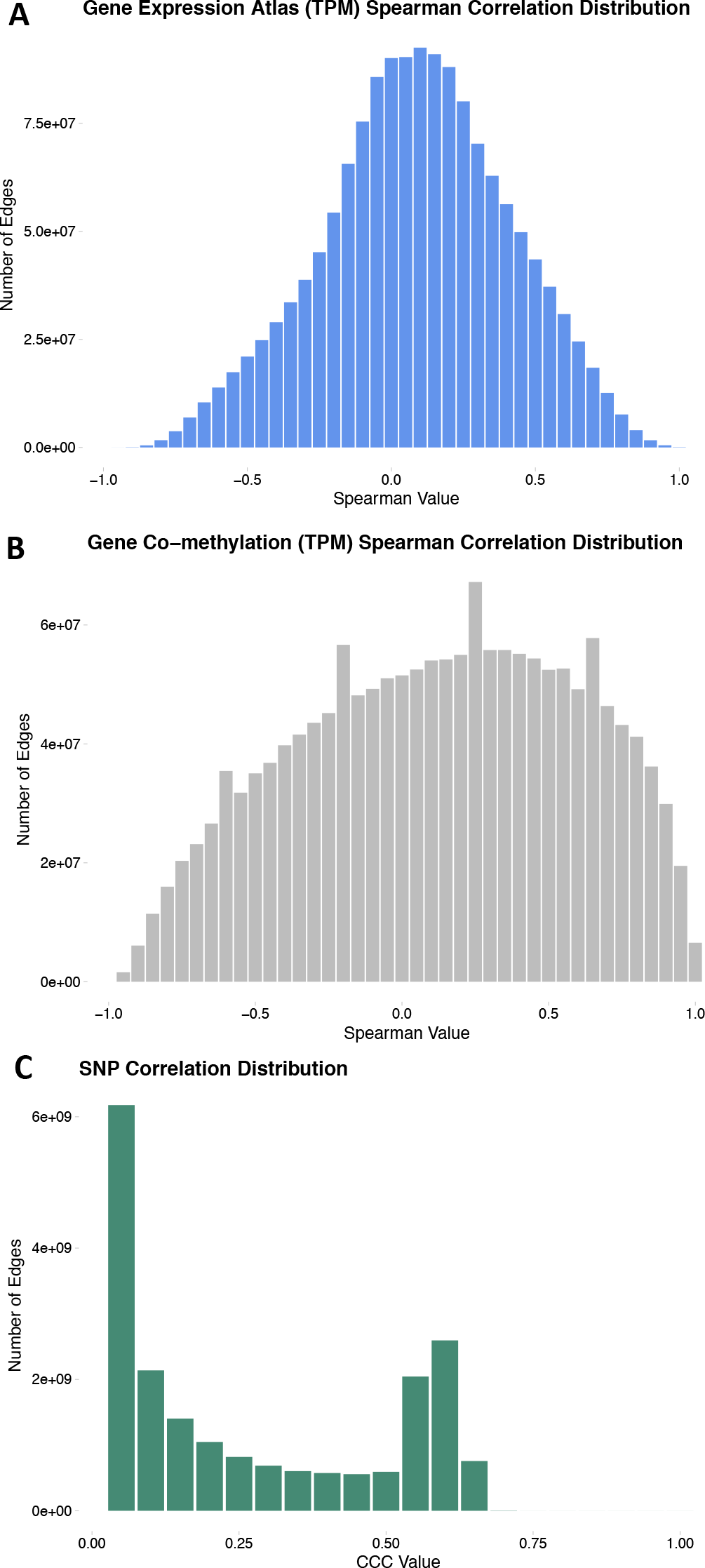
(A) Distribution of Spearman Correlation values in the co-expression network. (B) Distribution of Spearman Correlation values in the co-methylation network. (C) Sampled distribution of the CCC SNP correlation network. See Supplementary Note 1 for details on the construction of the sampled distribution.

**Figure S2:**
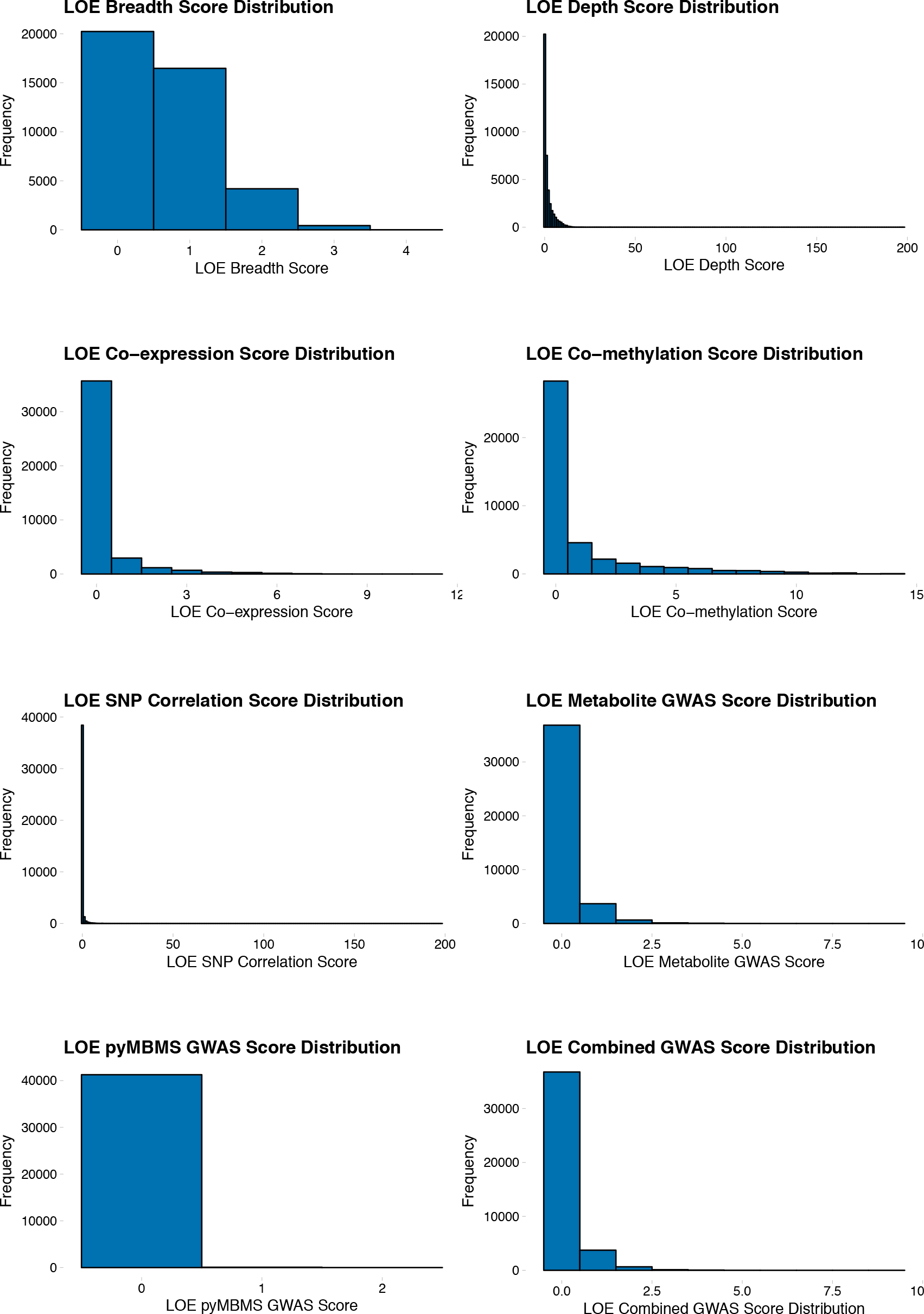
Lines Of Evidence (LOE) score distributions.

### 3. SUPPLEMENTARY TABLES

**Table S1:**
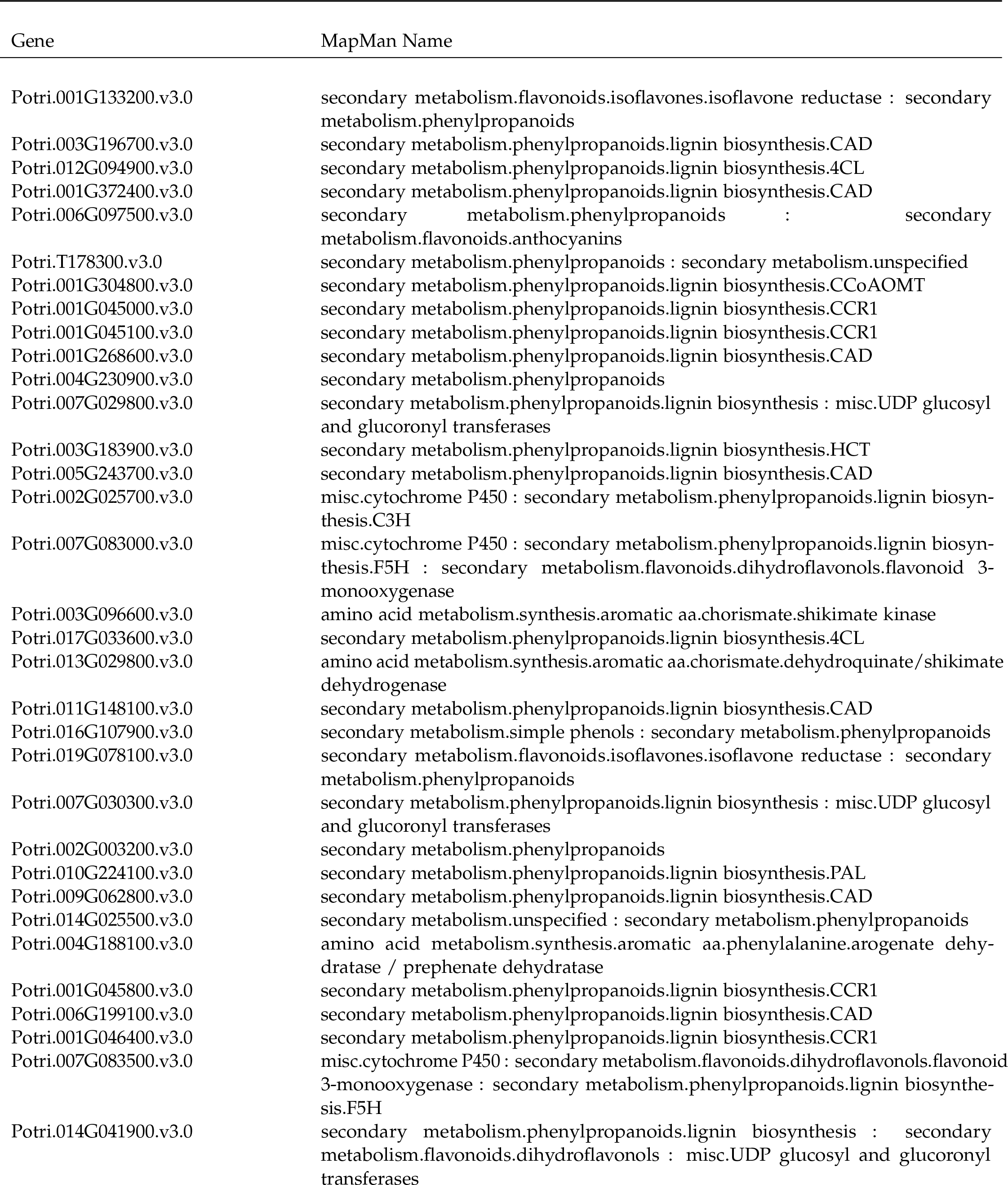
MapMan annotations of lignin genes.

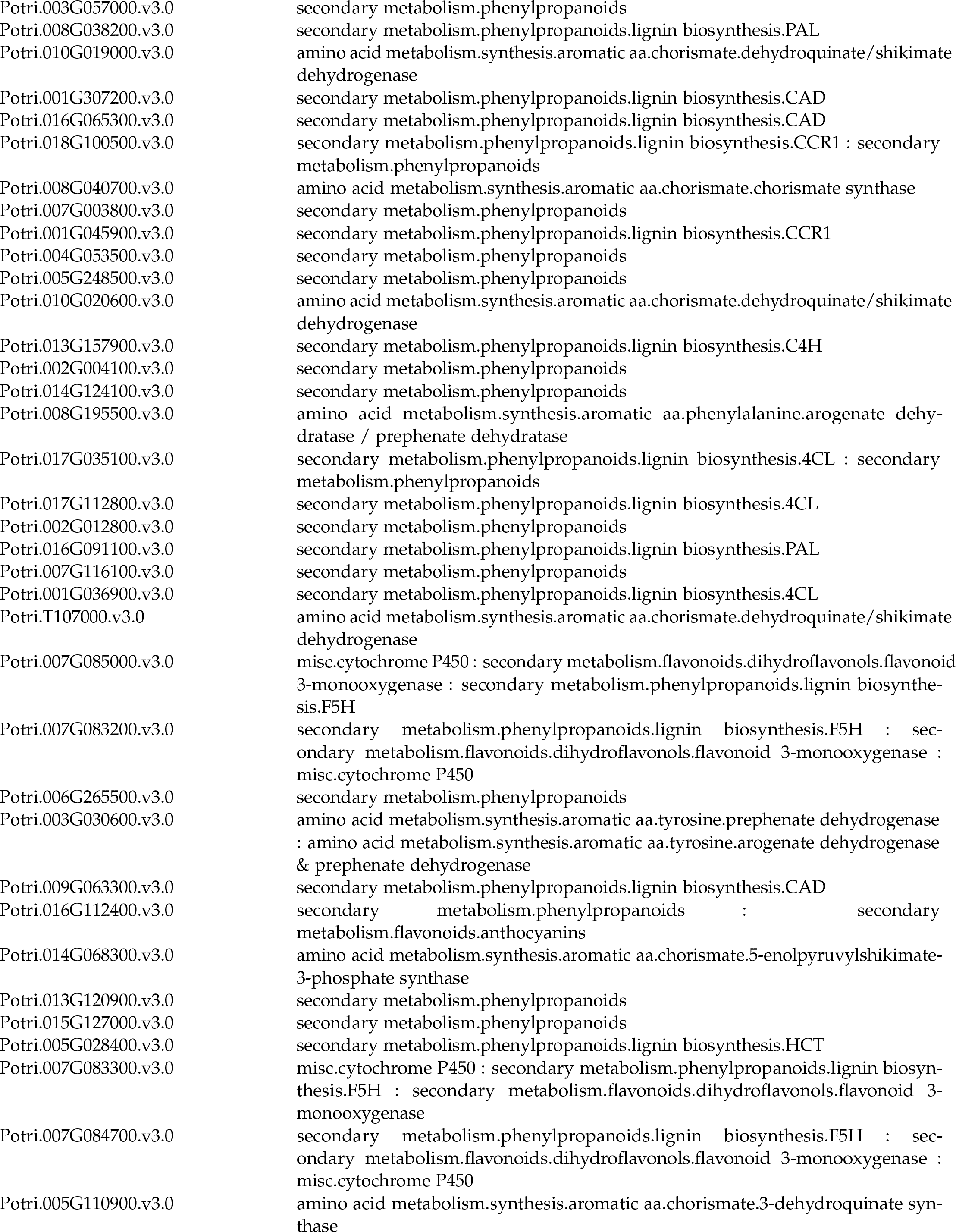

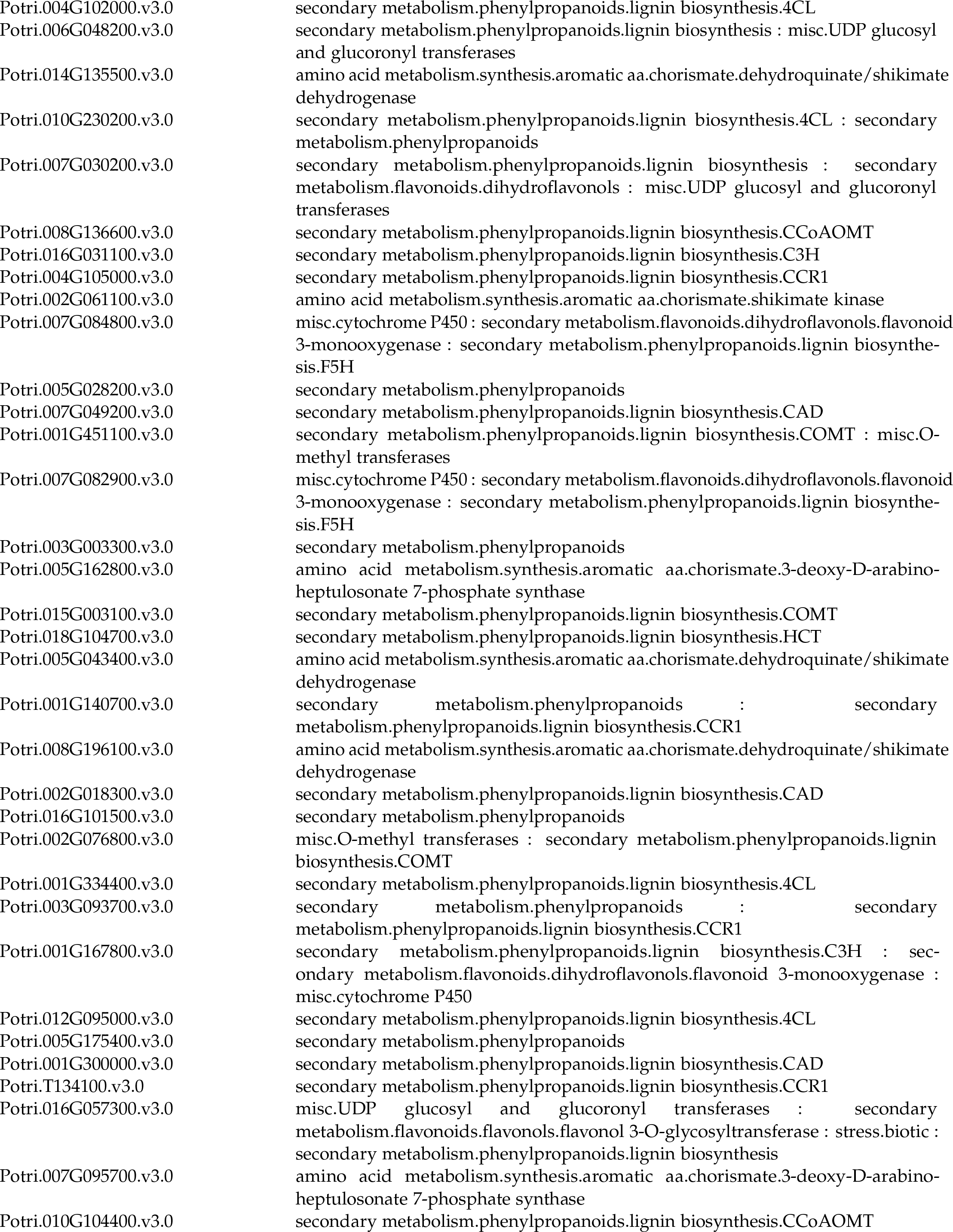

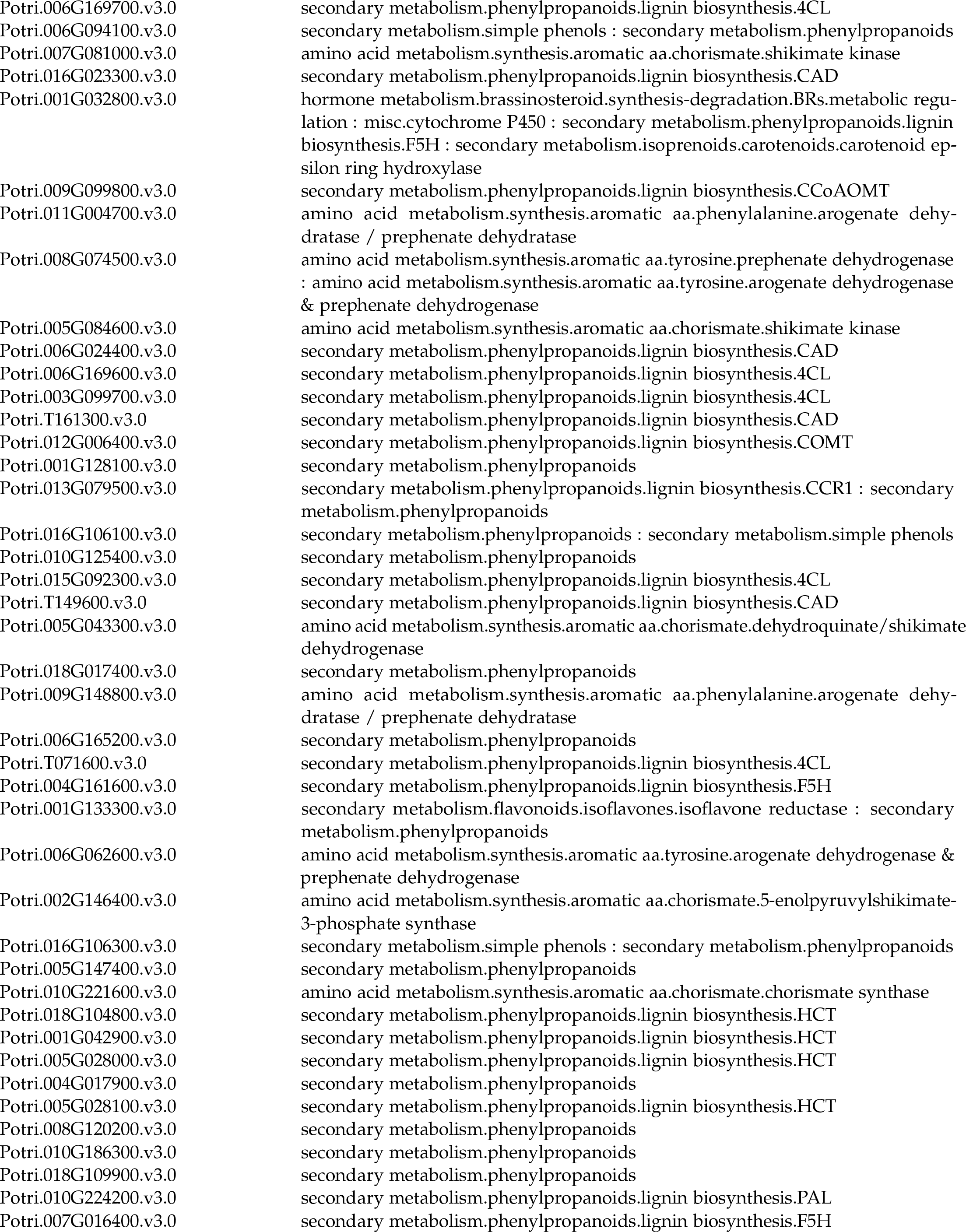

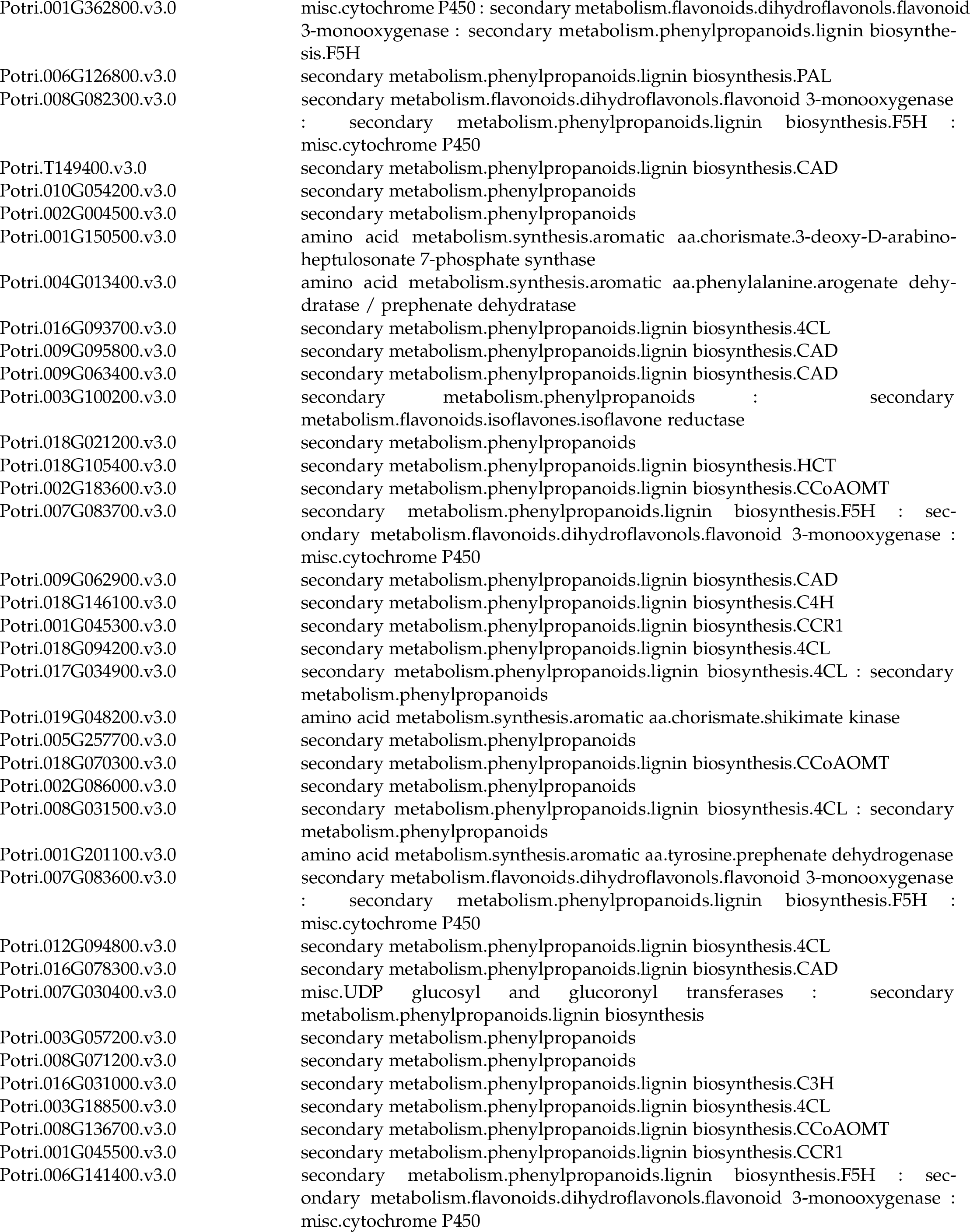

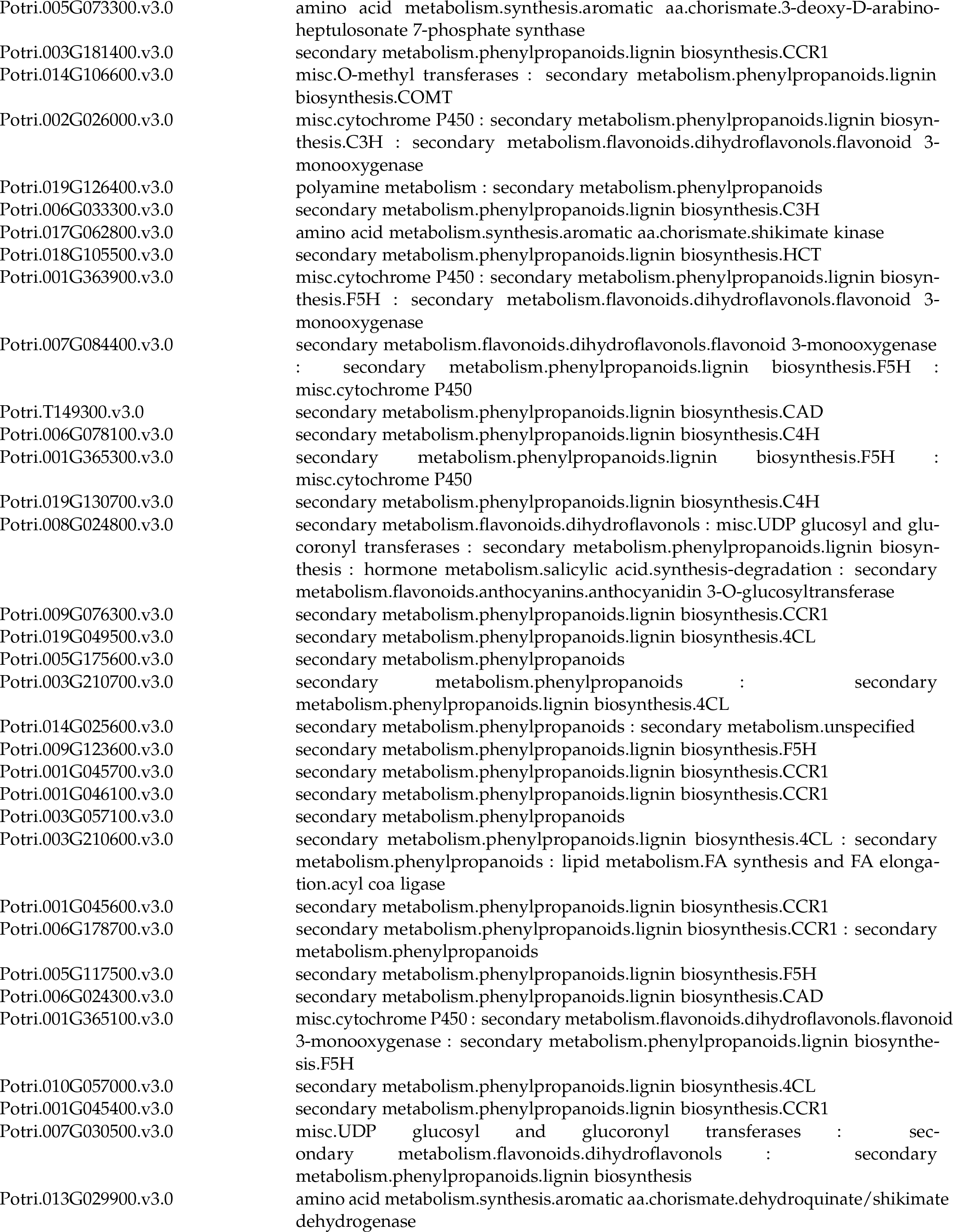

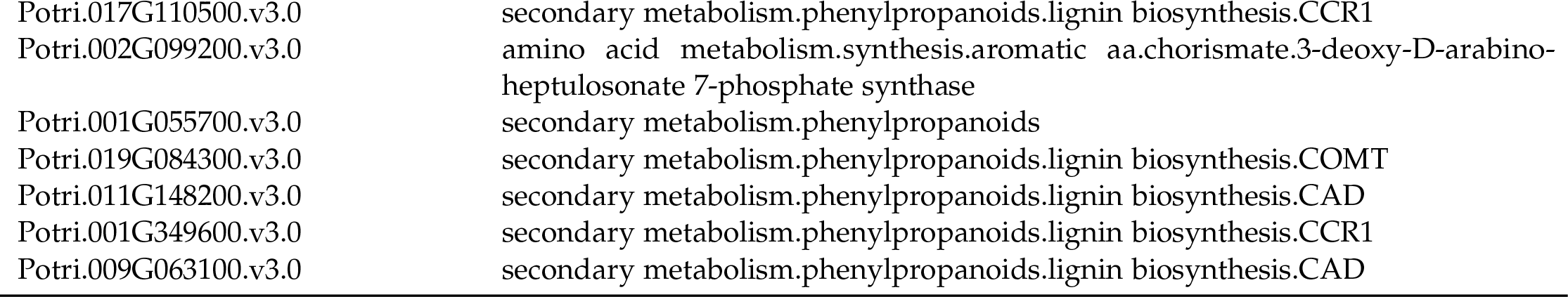

**Table S2:**
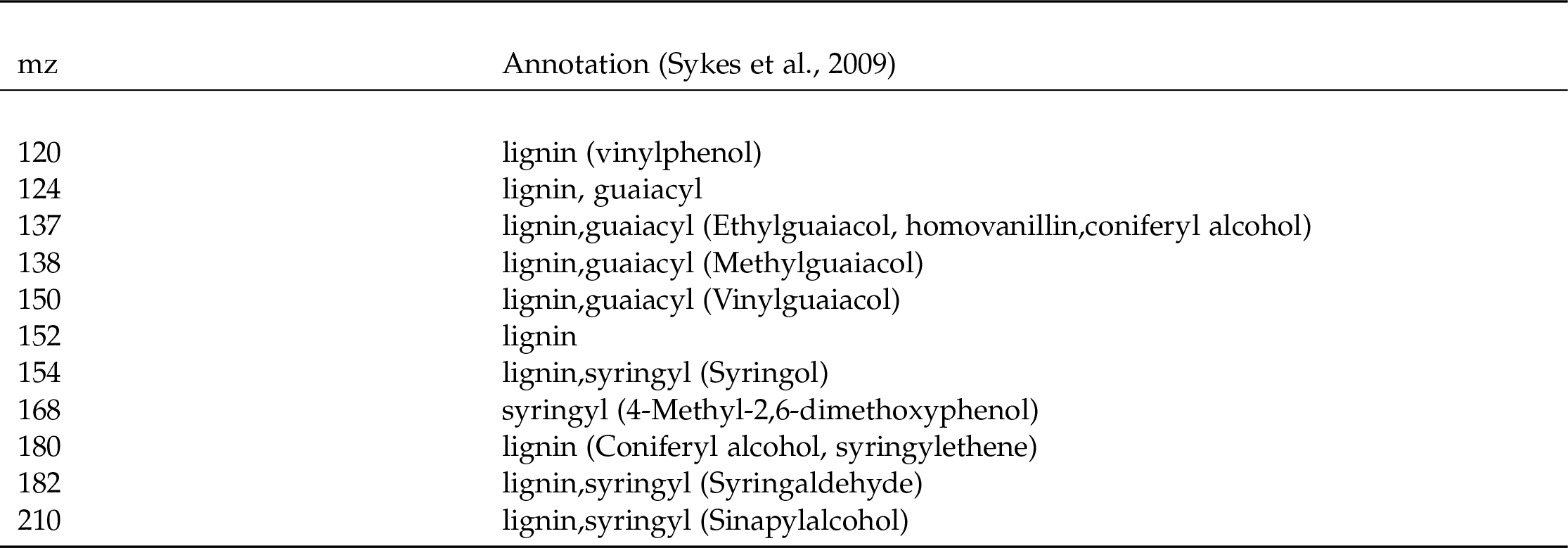
Mass/Charge (mz) ratio for Lignin pyMBMS Peaks.

| mz | Annotation (Sykes et al., 2009) |
| --- | --- |
| 120 | lignin (vinylphenol) |
| 124 | lignin, guaiacyl |
| 137 | lignin,guaiacyl (Ethylguaiacol, homovanillin,coniferyl alcohol) |
| 138 | lignin,guaiacyl (Methylguaiacol) |
| 150 | lignin,guaiacyl (Vinylguaiacol) |
| 152 | lignin |
| 154 | lignin,syringyl (Syringol) |
| 168 | syringyl (4-Methyl-2,6-dimethoxyphenol) |
| 180 | lignin (Coniferyl alcohol, syringylethene) |
| 182 | lignin,syringyl (Syringaldehyde) |
| 210 | lignin,syringyl (Sinapylalcohol) |

**Table S3:**
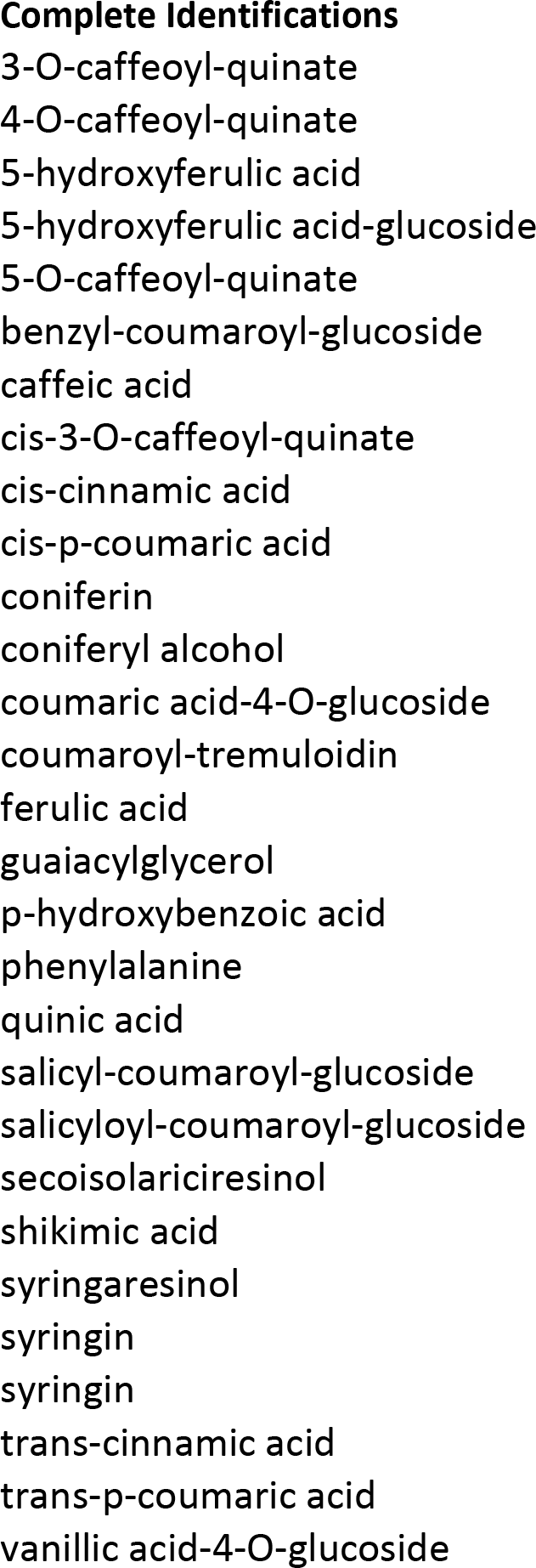
Lignin-related metabololites from the metabolomics analysis. For partially identified metabolites, additional RT and mz information is provided.

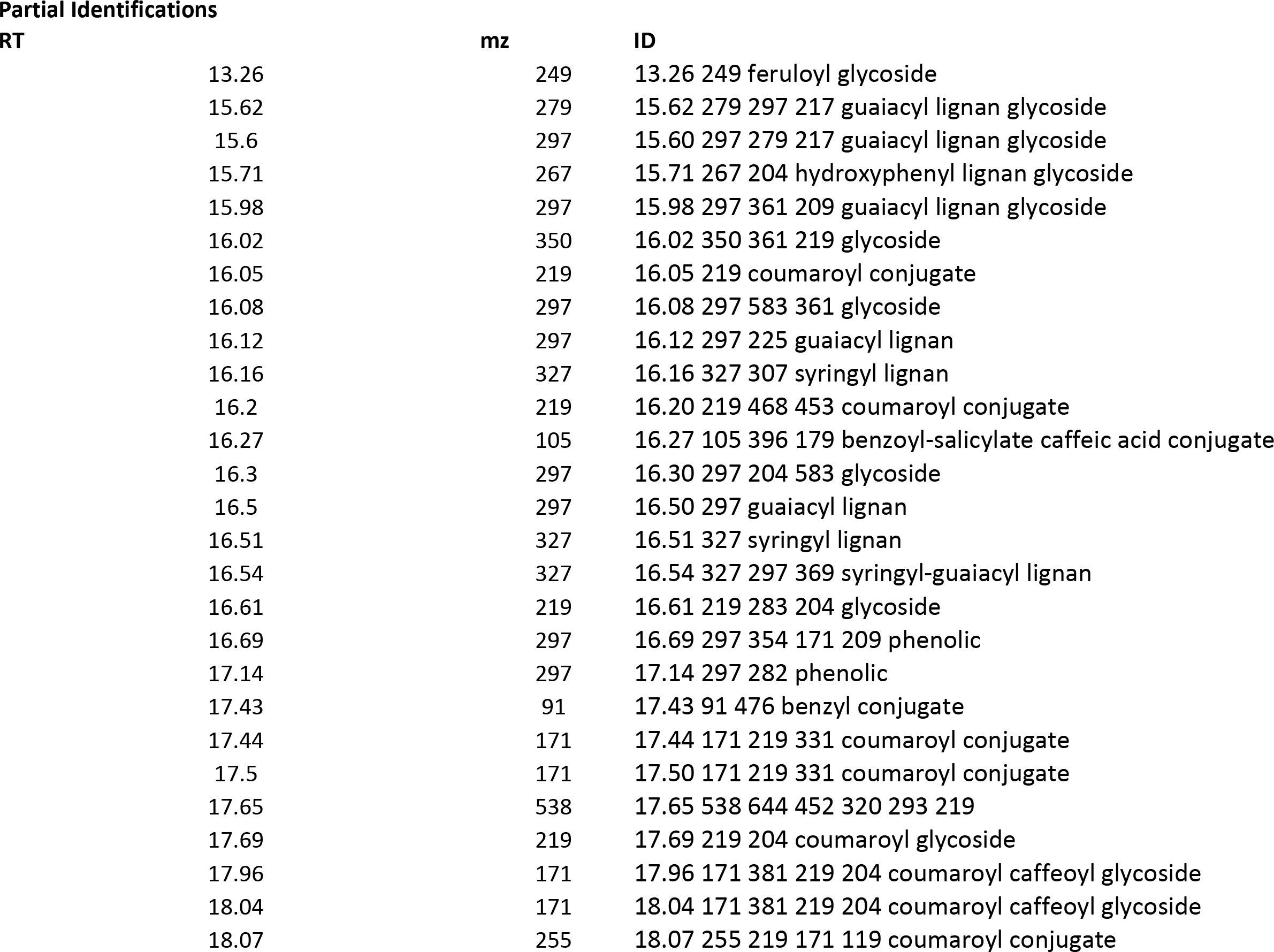

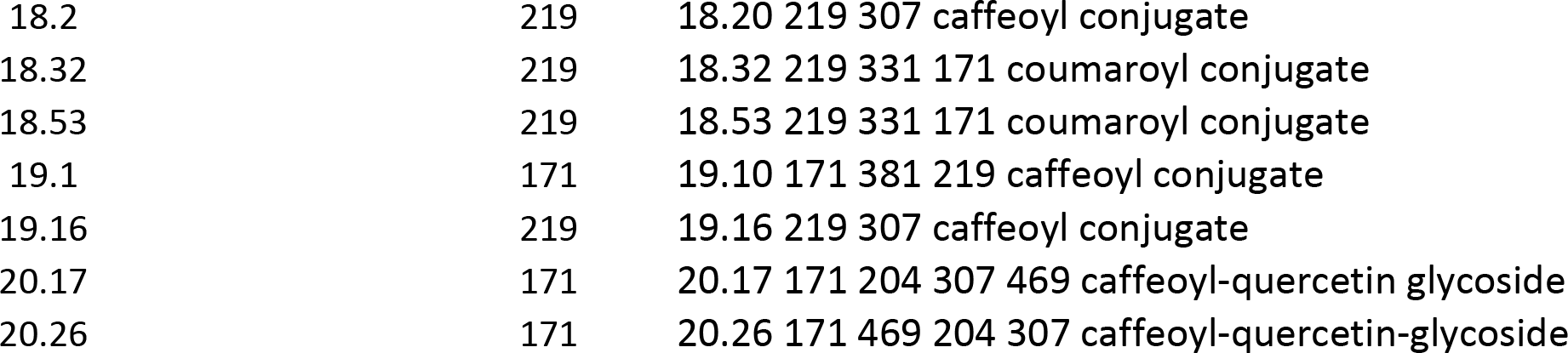

**Table S4:**
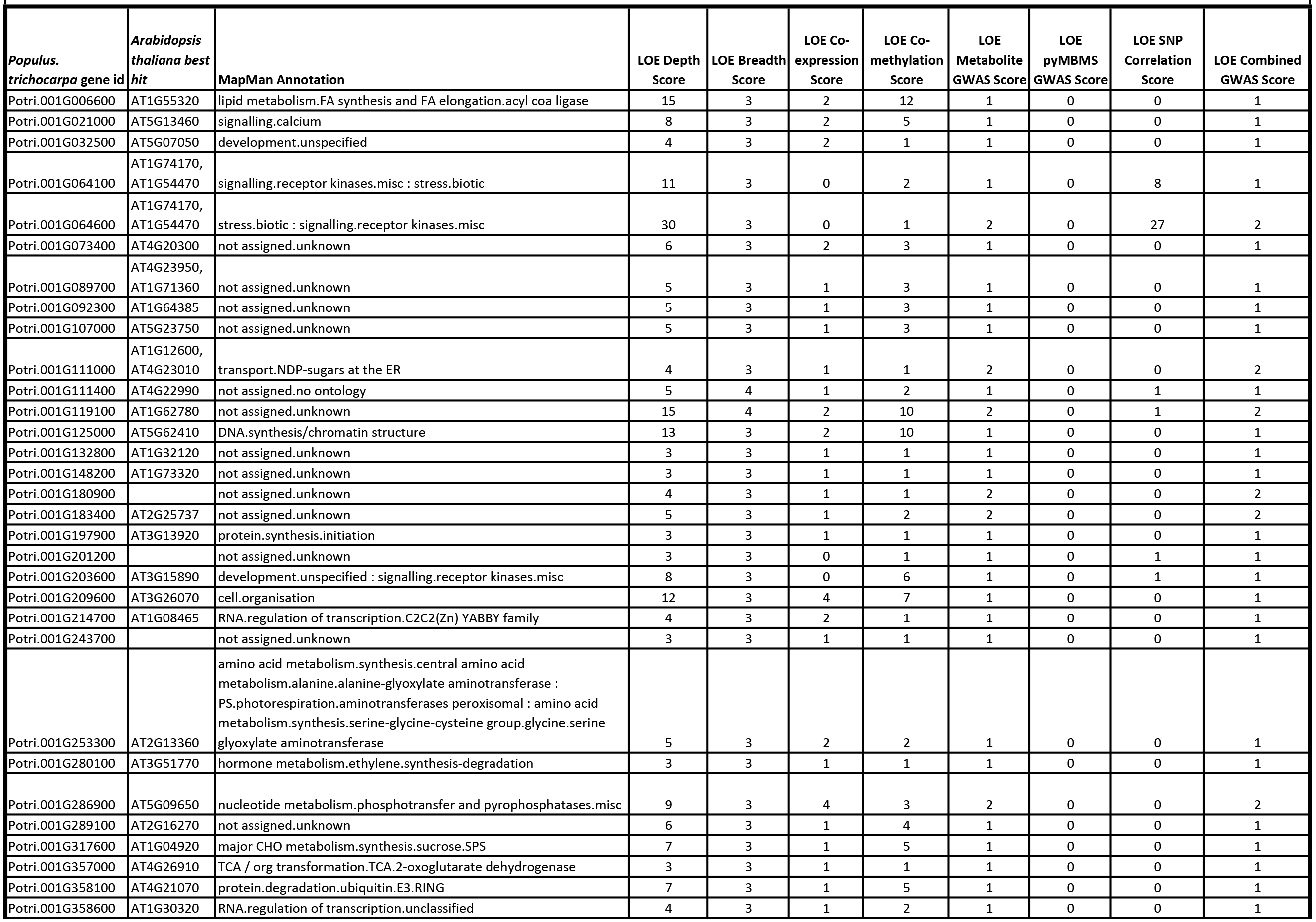
LOE Scores, Arabidopsis best hits and MapMan annotations of genes for which LOE_breadth_(g) ≥ 3 and LOE_gwas_(g) ≥ 1.

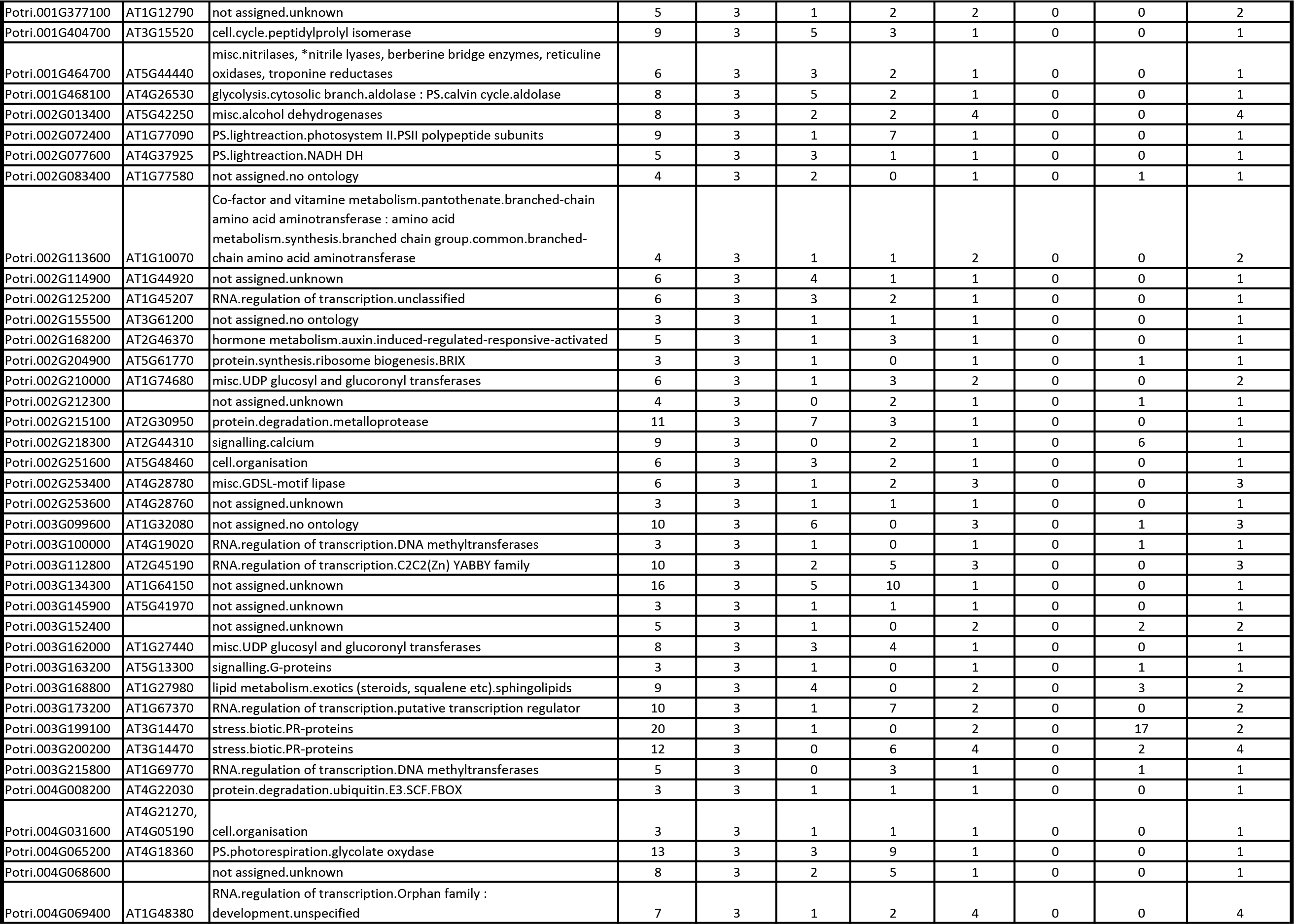

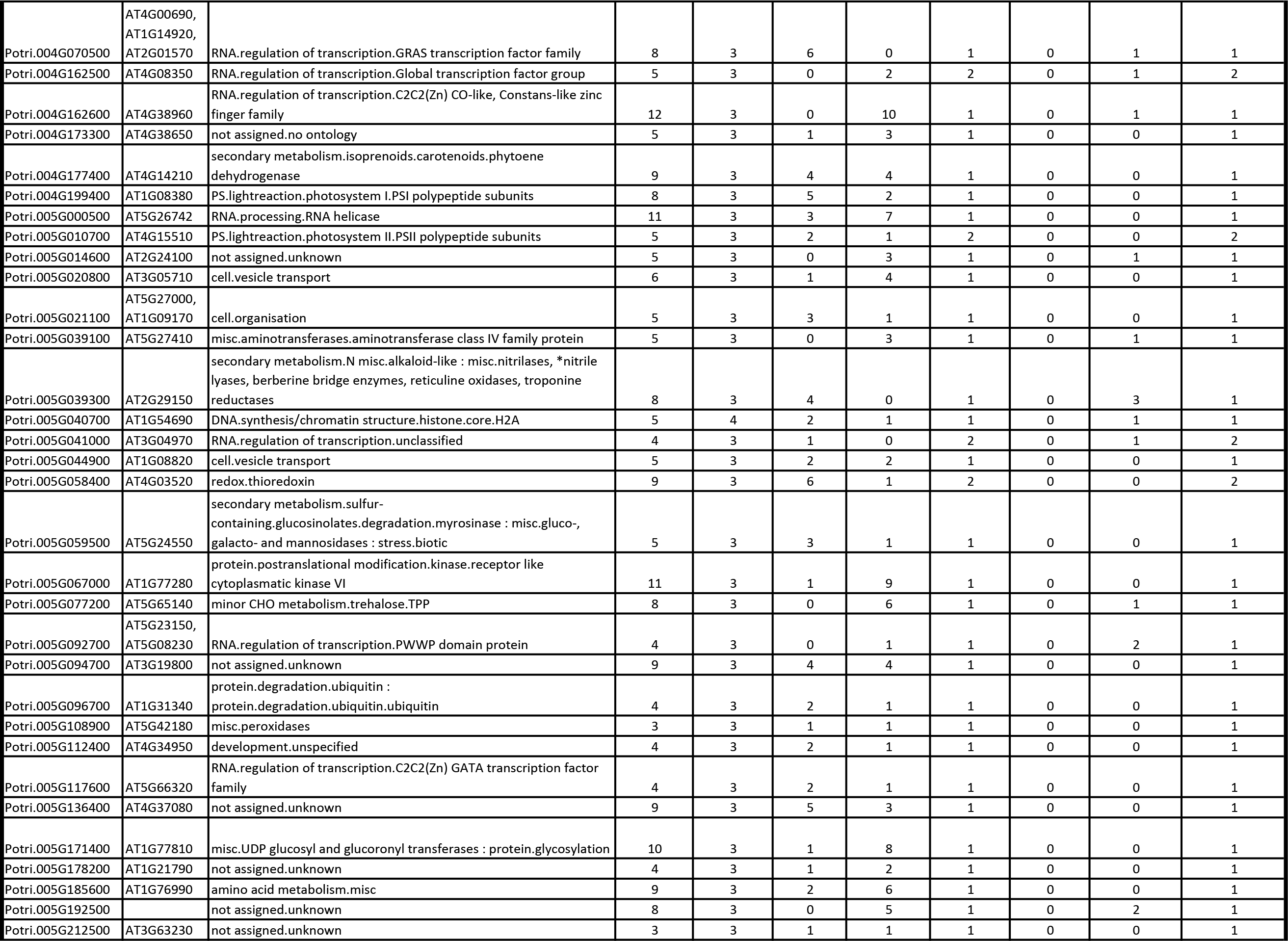

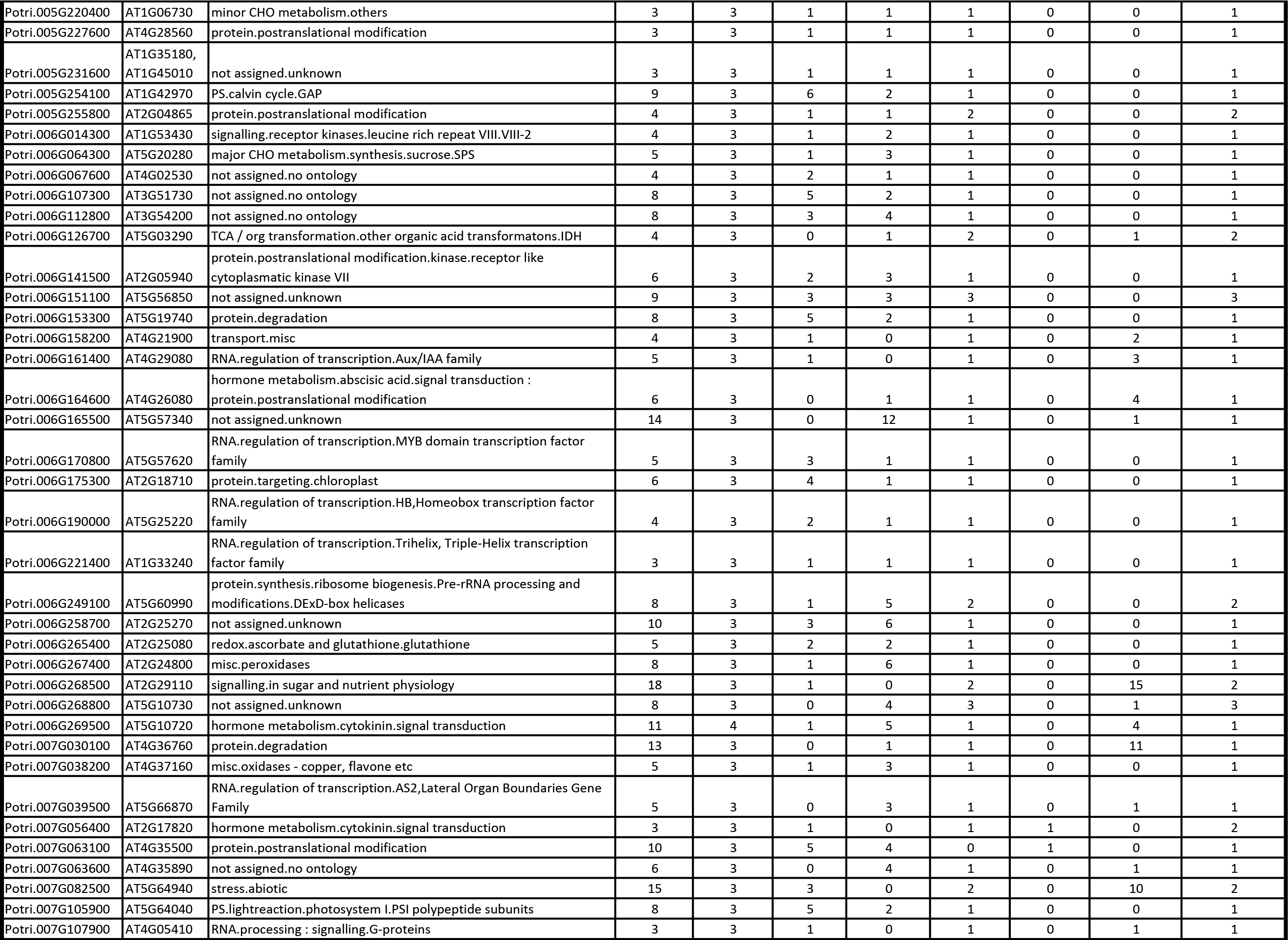

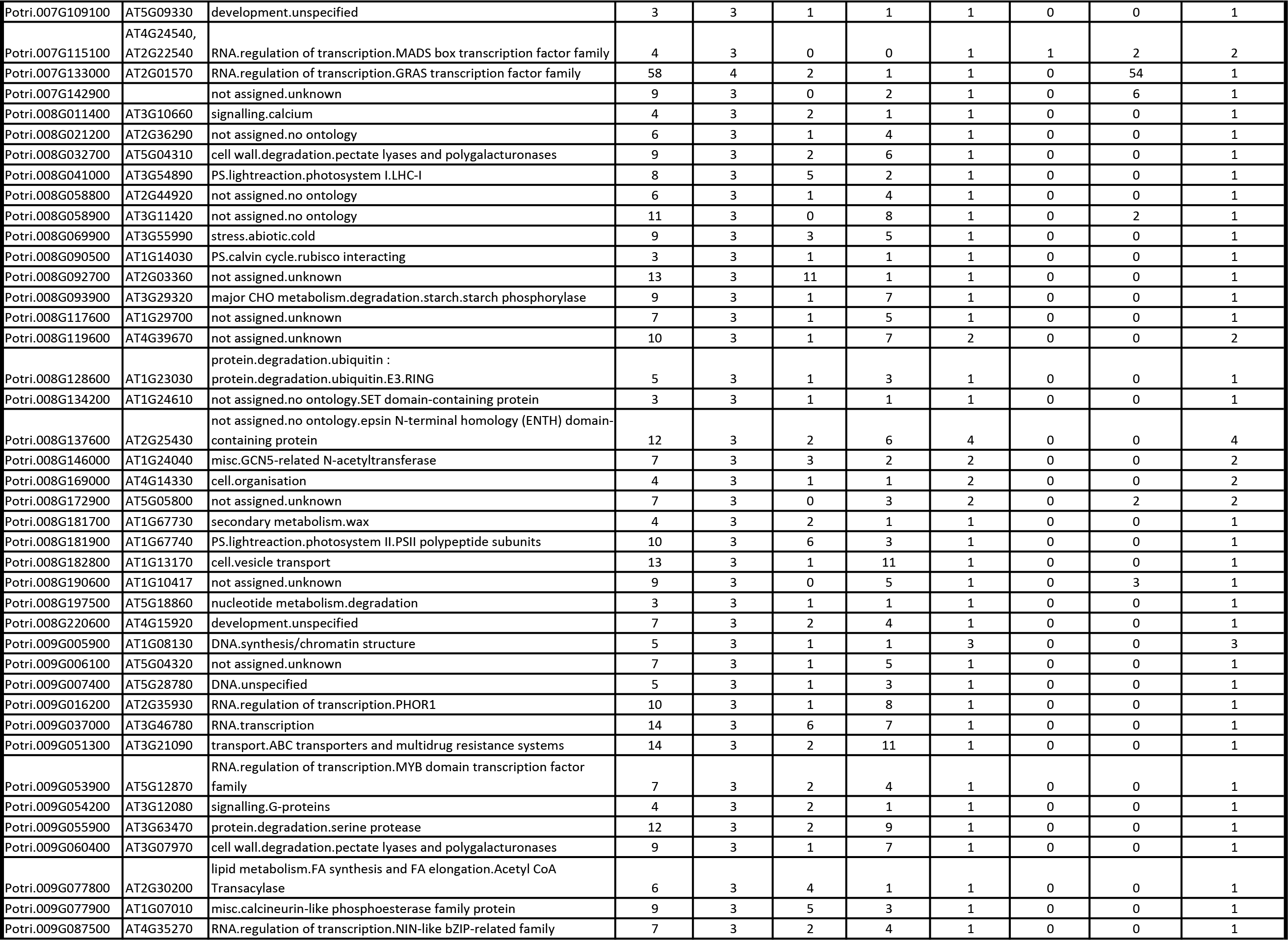

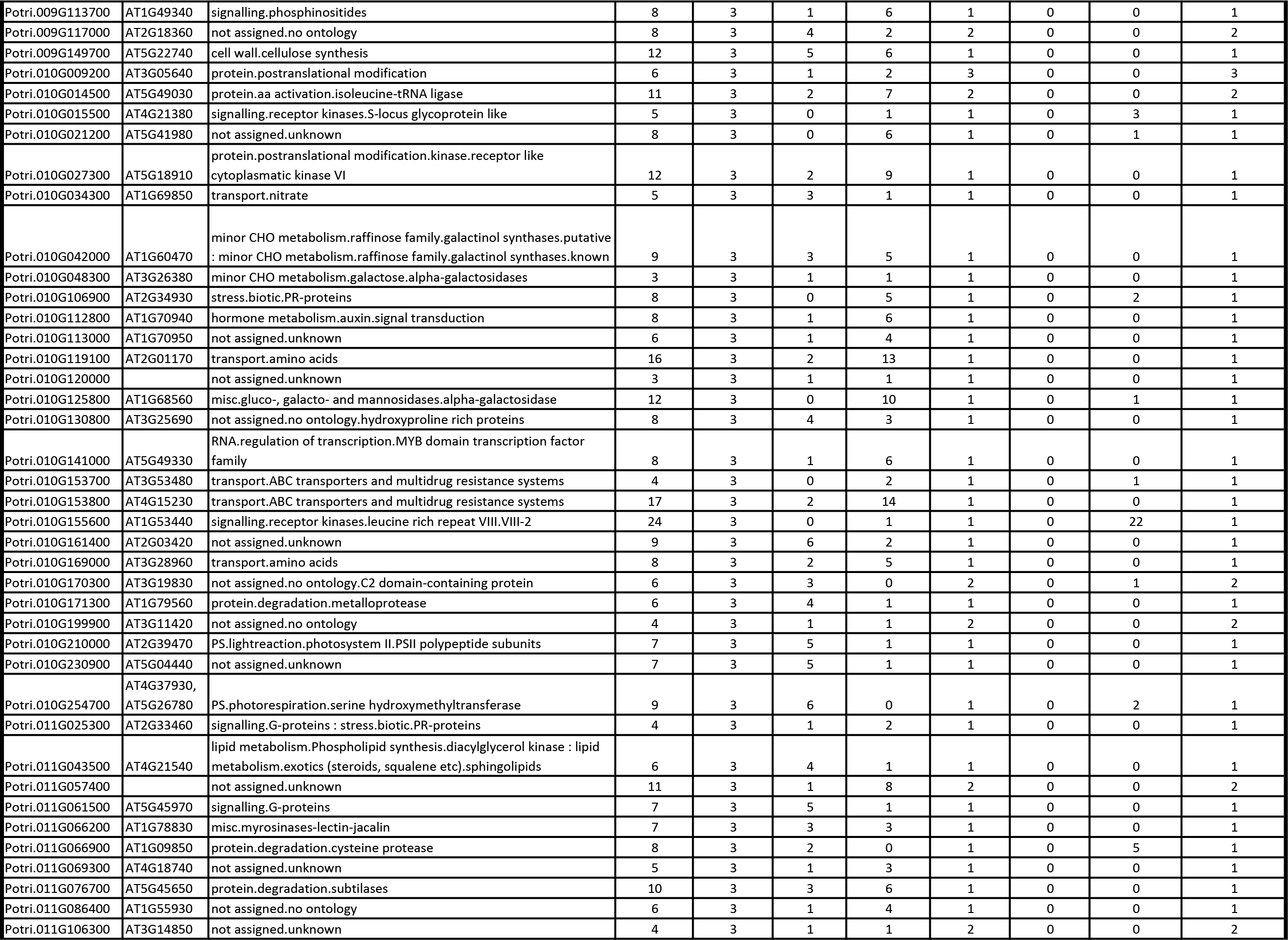

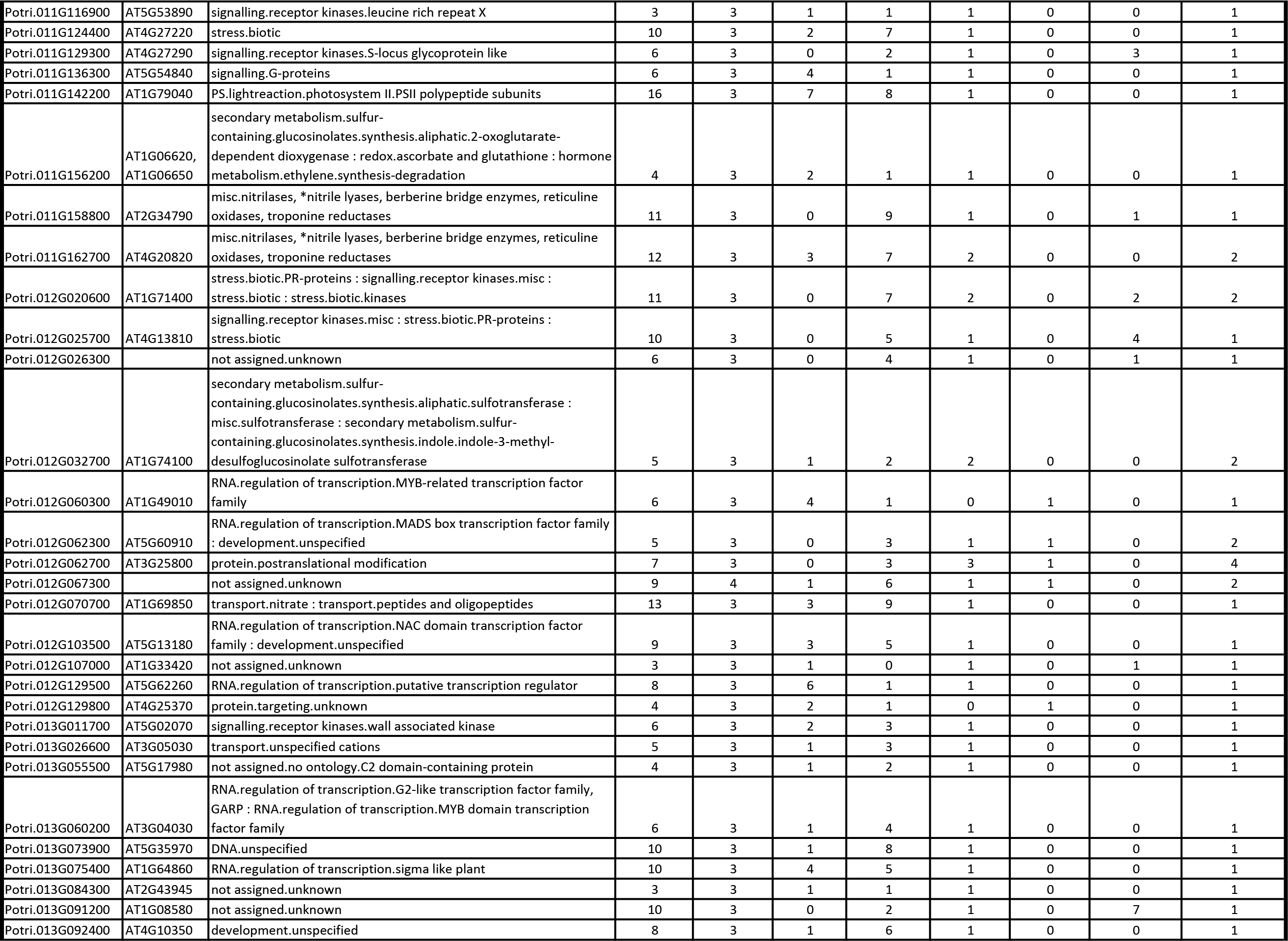

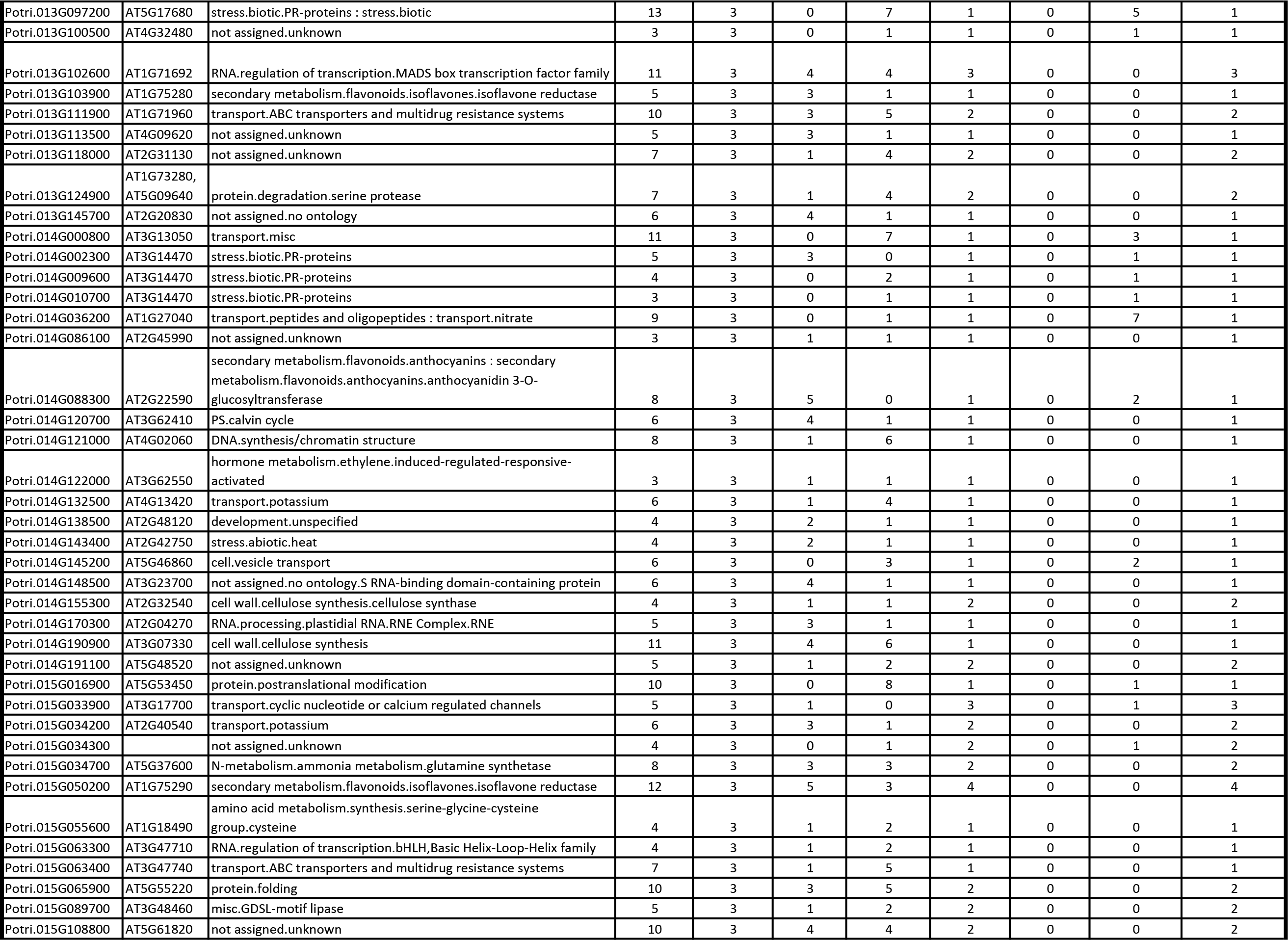

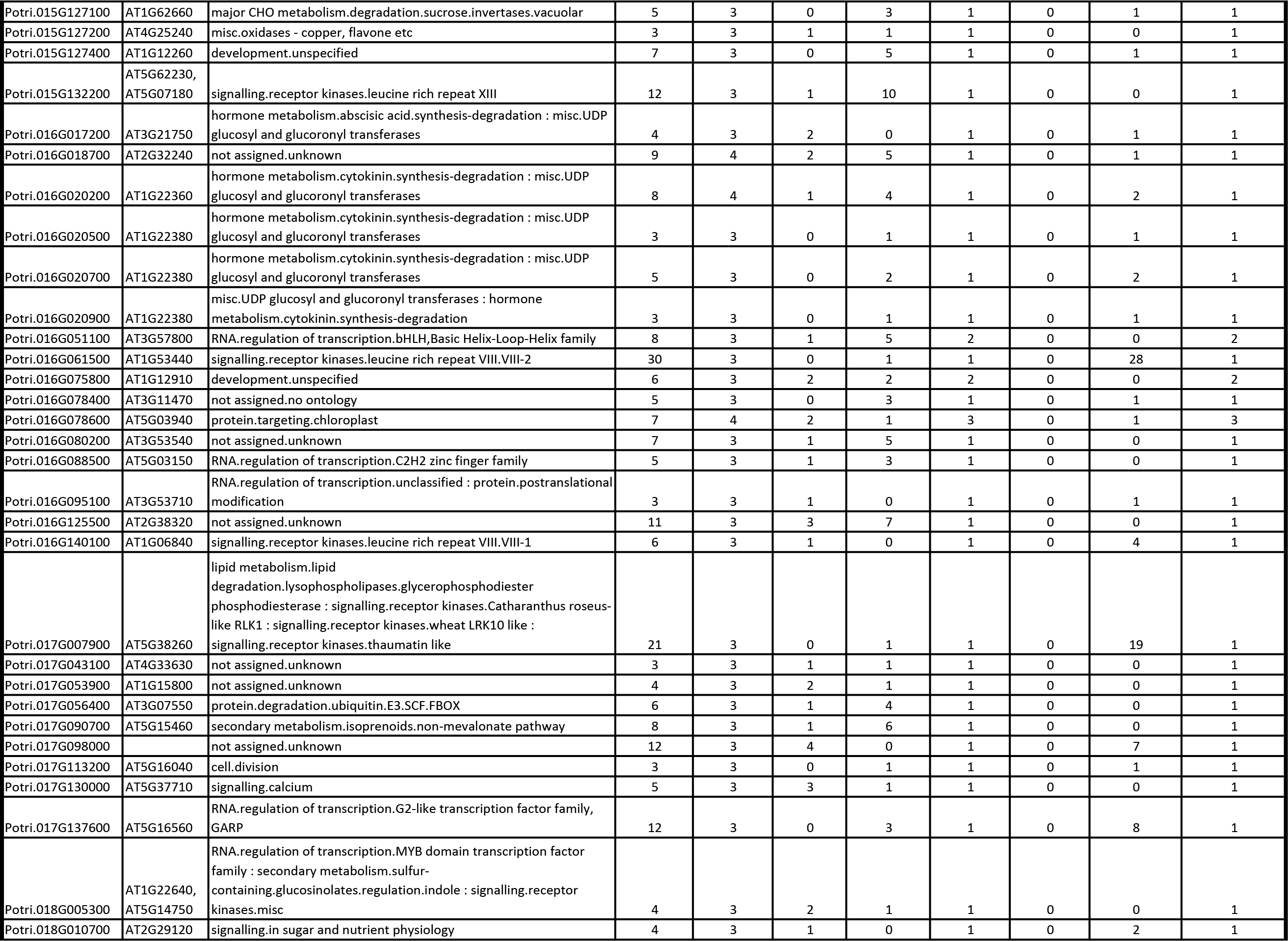

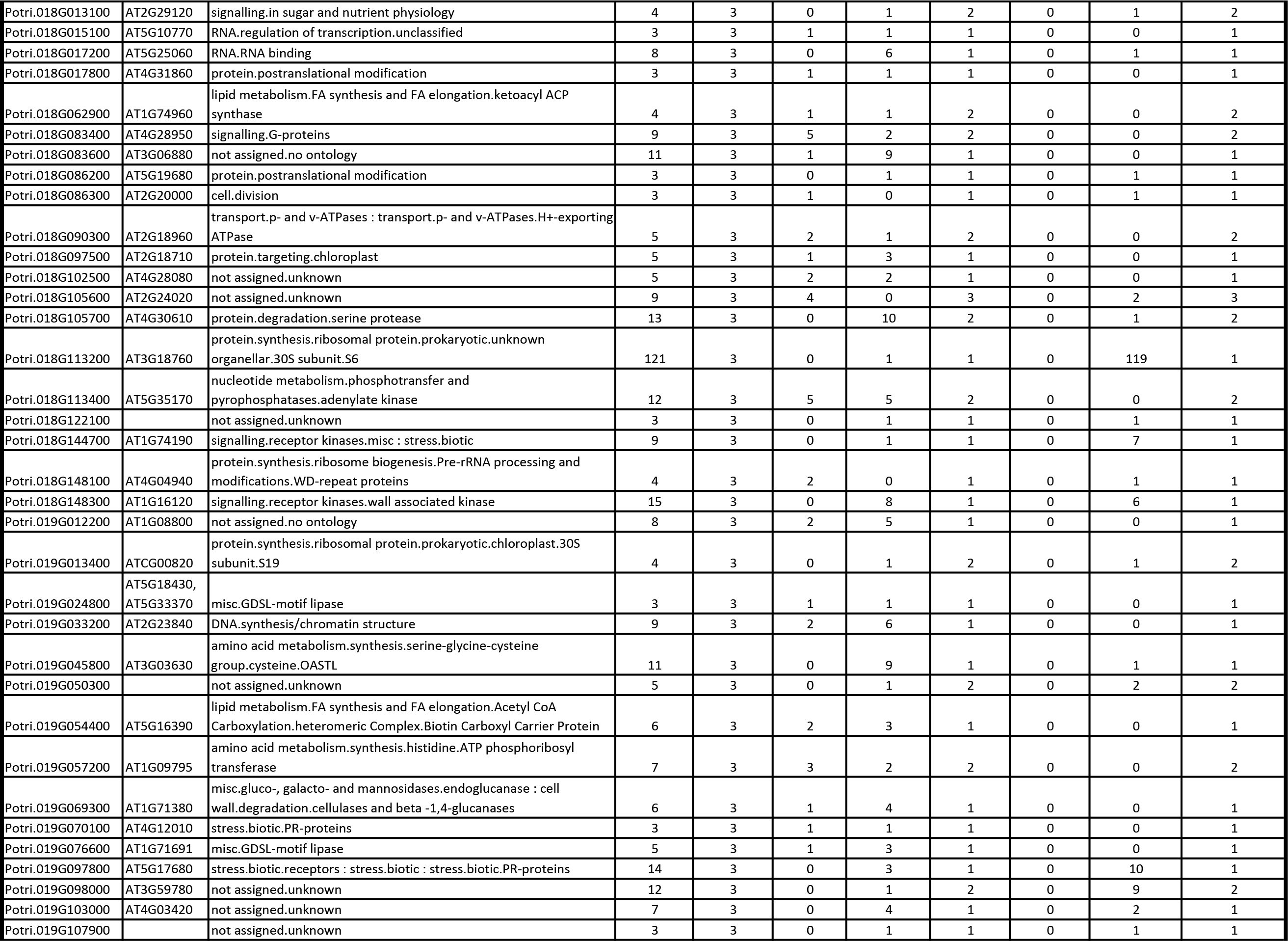

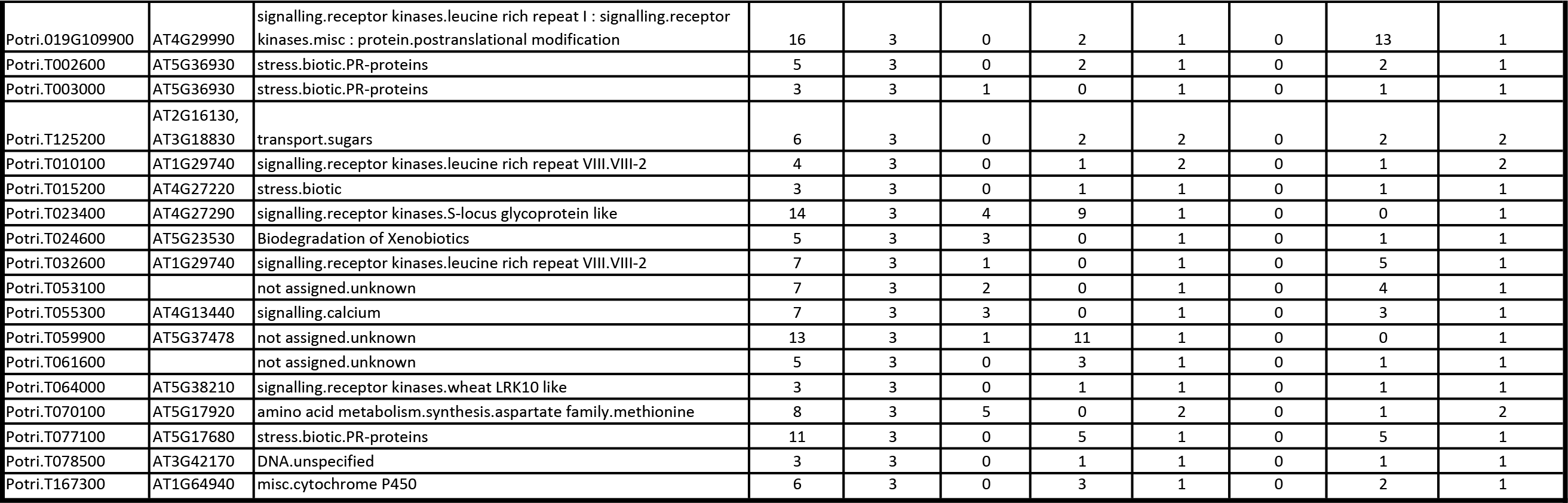

**Table S5:** Positions of SNPs involved in SNP correlations in select portential new target genes.

| Source | Target |
| --- | --- |
| **SNP Correlations between Potri.007G115100 (AGL22) and Potri.016G107900 (laccase 7)** |  |
| SNP 07:13647758 | SNP 16:11083690 |
| SNP 07:13647758 | SNP 16:11083708 |
| SNP 07:13647758 | SNP 16:11083712 |
| SNP 07:13647758 | SNP 16:11083737 |
| SNP 07:13647978 | SNP 16:11083690 |
| SNP 07:13647978 | SNP 16:11083708 |
| SNP 07:13647978 | SNP 16:11083712 |
| SNP 07:13647978 | SNP 16:11083737 |
| SNP 07:13648235 | SNP 16:11083690 |
| SNP 07:13648235 | SNP 16:11083708 |
| SNP 07:13648235 | SNP 16:11083712 |
| SNP 07:13648235 | SNP 16:11083737 |
| SNP 07:13648488 | SNP 16:11083690 |
| SNP 07:13648488 | SNP 16:11083708 |
| SNP 07:13648488 | SNP 16:11083712 |
| SNP 07:13648488 | SNP 16:11083737 |
| **SNP Correlations between Potri.007G115100 (AGL22) and Potri.007G116100 (nicotinamidase 1)** |  |
| SNP 07:13647645 | SNP 07:13706654 |
| SNP 07:13647645 | SNP 07:13706699 |
| SNP 07:13647645 | SNP 07:13706834 |
| SNP 07:13647758 | SNP 07:13706654 |
| SNP 07:13647758 | SNP 07:13706699 |
| SNP 07:13647978 | SNP 07:13706654 |
| SNP 07:13647978 | SNP 07:13706699 |
| SNP 07:13647978 | SNP 07:13706834 |

## Notes

This manuscript has been authored by UT-Battelle, LLC under Contract No. DE-AC05-00GR22725 with the U.S. Department of Energy. The United States Government retains and the publisher, by accepting the article for publication, acknowledges that the United States Government retains a non-exclusive, paid-up, irrevocable, worldwide license to publish or reproduce the published form of this manuscript, or allow others to do so, for United States Government purposes. The Department of Energy will provide public access to these results of federally sponsored research in accordance with the DGE Public Access Plan (http://energy.gov/downloads/doe-public-access-plan). The work conducted by the U.S. Department of Energy Joint Genome Institute is supported by the Office of Science of the U.S. Department of Energy under Contract No. DE-AC02-05CH11231.

